# mRNA isoform switching plays a crucial role in mural cumulus differentiation

**DOI:** 10.1101/2025.11.11.684658

**Authors:** Sharmin Shila, Grace J. Pei, Niki Peramsetty, Elizabeth Bahadursingh, Vinesh Dahiya, Courtney A. Marsh, Ramkumar Thiyagarajan, Meijia Zhang, Patrick E. Fields, M. A. Karim Rumi

**Affiliations:** Department of Pathology and Laboratory Medicine, University of Kansas Medical Center, Kansas City, KS, USA; Department of Obstetrics and Gynecology, University of Kansas Medical Center, Kansas City, KS, USA; Department of Internal Medicine, University of Kansas Medical Center, Kansas City, KS, USA; Division of Cell, Developmental and Integrative Biology, South China University of Technology, School of Medicine, Guangzhou, China

**Keywords:** gonadotropin response, antral follicle, mural granulosa cells, cumulus granulosa cells, transcriptome analysis, transcript isoforms, transcript switching

## Abstract

During gonadotropin-induced ovarian follicle development, an antral cavity forms within the layers of granulosa cells (GCs). Gonadotropin stimulation also differentiates the GCs into two distinct lineages: mural GCs (mGCs), which surround the antral cavity, and cumulus GCs (cGCs), which stay in contact with the oocyte. We examined the transcriptomes of mouse mural and cumulus cells to understand the mechanism of differentiation. In addition to analyzing a single transcript expression per gene, we also considered multiple isoform expressions to explore differential transcriptomics. GC-specific core transcription factors *Foxl2, Nr5a1, Nr5a2, Runx1*, and *Runx2* were expressed at high levels in mGCs but downregulated in cGCs, indicating that cGCs acquire a more differentiated state. Both single-transcript and multiple-isoform analyses revealed differential expression of about 70% of transcripts between mGCs and cGCs. Although the counts were similar, the differentially expressed genes (DEGs) at the single transcript level did not correlate well with the respective differentially expressed transcript isoforms (DETI). We identified DETIs originating from key epigenetic and transcriptional regulator genes, such as *Chd1, Ezh2, Kdm5a, Kdm5b, Gata4, Esr2, Fos, Myc*, and *Ybx1*, that were not differentially expressed at the single-transcript analysis. Further analysis revealed a transcript switch in one-third of the DETIs. Most of the transcript isoforms were protein-coding, followed by non-coding regulatory RNAs. A total of 1,302 transcript isoforms were silenced in cGCs, including those of Adar, Cebpa, *Dnmt3a, Foxo4, Pgr, Rest, Runx1, Satb2, Sirt1, Sirt2*, and *Tead1*. Conversely, 529 transcript isoforms were activated in cGCs, including transcripts for *Brd7, Crem, Chd1, Med21, Med27, Nfkbia, Rbm39, Rbmx, Suv39h2, Tcf12, Xist*, and *Ybx3*. Additionally, 57 genes exhibited DETIs, with at least one isoform turned off and another turned on in cGCs, including *Csde1, Dab2, Ezh2, Gata4, Gnas, Gtf2i, Macf1, Klf10, Setdb1*, and *Sp3*. Finally, we explored the mechanisms underlying transcript switching during the differentiation of mGCs and cGCs. Our findings suggest that gonadotropin-induced transcript switching in GCs is crucial for mural and cumulus granulosa differentiation, a key insight that would be missed without mRNA isoform analysis.

## 1. INTRODUCTION

Granulosa cells (GCs) continue to grow and differentiate throughout ovarian folliculogenesis (1). The somatic cells of preantral follicles express gonadotropin receptors, making follicle maturation responsive to gonadotropins (2). Two gonadotropins, follicle-stimulating hormone (FSH) and luteinizing hormone (LH), act on the GCs. During gonadotropin-induced follicle development, an antral cavity forms within the layers of GCs, and stimulation by gonadotropins causes the GCs to differentiate into two distinct populations (3, 4). Cumulus GCs (cGCs) remain in contact with the oocytes, while mural GCs (mGCs) surround the antral cavity near the basement membrane and theca cells (5, 6). Although mGCs mainly produce ovarian steroids, cGCs support oocyte maturation (7, 8). This structural and functional division ensures the follicle is fully prepared for ovulation. The antral cavity is a fluid-filled space containing secretions from GCs and oocyte-derived factors that facilitate nutrient and hormone exchange between GCs and oocytes, support oocyte maturation, and generate the pressure needed for follicle rupture during ovulation (9, 10).

mGCs and cGCs originate from GCs proper, but they show significant differences in gene expression. Even the gonadotropin receptors are expressed differently in these two types of GCs (11). Compared to mGCs, cGCs have higher levels of *Fshr* but lower levels of *Lhcgr* (12). In addition to gonadotropin receptors, many other genes are expressed differently between mGCs and cGCs. While steroidogenic enzymes *Cyp19a1, Cyp11a1*, and *Star* are upregulated in mGCs, glycolytic enzymes *Pgk1, Tpi*, and *Eno1* are upregulated in cGCs (13, 14). Due to their location and proximity, it is also suggested that gene expression in mGCs is regulated by theca cell-derived factors, while gene expression in cGCs is regulated by oocyte-derived factors (15, 16). We hypothesize that stimulation by theca or oocyte-derived factors influences lineage-specific epigenetic and transcriptional regulators, which are responsible for the differential gene expressions observed in mGCs and cGCs.

Studies often analyse gene expression to understand the mechanism of cell differentiation (17, 18). Several studies have demonstrated the differential expression of genes during the differentiation of mural and cumulus cells (19-22). However, such conventional gene expression analysis that considers a single mRNA expressed from a gene is biologically inaccurate (23). Most mammalian genes produce multiple transcript isoforms, and not all of these transcripts encode proteins (24). Some transcript isoforms are upregulated in precursor cells, while others are induced in differentiated cells (22). In this study, we have analyzed the transcriptomes of mouse mural and cumulus cells to investigate the mechanisms of differentiation. In addition to traditional single-mRNA (transcript) analysis, we also examined transcript isoform expression to identify differential transcriptomics. We found that many transcripts highly expressed in mGCs are repressed in cGCs. Conversely, a group of isoforms repressed in mGCs is induced in cGCs. Our results suggest that gonadotropin-induced transcript switching in GCs plays an essential role in the differentiation of mural and cumulus cells.

## Materials and Methods

### 2.1. Gonadotropin-induced granulosa cell differentiation

Ovaries in 3-week-old (21-23 days) C57BL/6 mice were stimulated with exogenous gonadotropins to induce granulosa cell proliferation and differentiation. Mice received an intraperitoneal injection of 5 IU of eCG. Forty-eight hours later, the eCG-primed mice were injected intraperitoneally with 5 IU of hCG (25, 26). Two hours after the hCG injection, the mice were euthanized, and their ovaries were collected. For histological analysis, the ovaries were fixed in 4% formaldehyde overnight, processed, and embedded in paraffin following standard procedures (27, 28). Whole ovaries were serially sectioned at 6 µm thickness and stained with hematoxylin and eosin. All animal experiments were approved by the Institutional Animal Care and Use Committee of South China University of Technology (25) and the University of Kansas Medical Center (29).

### 2.2. Isolation of mural and cumulus cells and purification of RNA

mGCs and cGCs were isolated from gonadotropin-stimulated mouse ovaries (25). Mural cells and cumulus oocyte complexes (COCs) were expressed from the antral follicles using needle picking under stereoscopic examination. The COCs were transferred to a different plate with a pipette, and cumulus cells were separated from the oocytes by repeated pipetting (30, 31). Both types of GCs were purified using strainers, washed with cold PBS, and mGCs were purified from RBCs using a standard lysis procedure. Isolated mural and cumulus cells were used for RNA extraction and RNA-sequencing library preparation, sequencing, and other downstream analyses (25, 32, 33).

### 2.3. RNA sequencing

Total RNAs purified from mGCs and cGCs with high RIN values were used for bulk RNA-sequencing (25). The libraries were sequenced on an Illumina NovaSeq 6000 platform. The data were generated by the Zhang laboratory at South China University of Technology, Guangzhou, China, and have been submitted to the SRA, NCBI (PRJNA891667). The datsets included three mGCs transcriptomes (SRR21944927, SRR21944928, and SRR21944929) and three cGCs transcriptomes (SRR21944921, SRR21944922, and SRR21944923). The RNA-seq data were analyzed and validated as described in the following sections.

### 2.4. Preparation of cDNAs and RT-PCR

Total RNA was extracted from the isolated mGCs or cGCs using TRIReagent (Millipore-Sigma). 1000 ng of total RNA from each sample was used to synthesize cDNA using the High-Capacity Reverse Transcription Kit (Applied Biosystems, Foster City, CA). The cDNA was diluted 1:25 in 10 mM Tris-HCl (pH 7.4), and 2.5 µL of this diluted cDNA was added to a 10-µL qPCR reaction mixture containing Applied Biosystems Power SYBR Green PCR Master Mix (ThermoFisher Scientific) (34). All PCR primers were designed using Primer3 (ver. 4.0) (35), and the sequences are shown in **Supplementary Table S1**. Amplification and fluorescence detection for RT-qPCR were performed using the Applied Biosystems QuantStudio Flex 7 Real-Time PCR System (Thermo Fisher Scientific). The ΔΔCT method was used for relative quantification of target mRNA normalized to 18S RNA (36).

### 2.5. RNA sequencing analysis

RNA-Seq data were processed using CLC Genomics Workbench 25.0.3 (Qiagen Bioinformatics, Redwood City, CA, USA) (37). All clean reads were produced by removing low-quality sequences and trimming adapter regions (38). The high-quality reads were aligned to the Mus musculus reference genome (GRCm39), gene (GRCm39.114_Gene), and mRNA sequences (GRCm39.114_mRNA) using default parameters. The expression levels of single transcripts (GE) or multiple transcript isoforms (TE) in mGC and cGC cells were measured in TPM. The p-value threshold was determined based on the false discovery rate (FDR) (39, 40). FDR p-value ≤0.05 was considered significant.

### 2.6. Detection of differentially expressed genes

Gene expression, whether based on a single transcript or multiple transcript isoforms, was analyzed only for those with ≥ 1 TPM. Differentially expressed genes were divided into three groups: upregulated (≥2-fold change and FDR p ≤0.05), downregulated (≤ -2-fold change and FDR p ≤0.05), and insignificant (either <2-fold change and/or FDR p >0.05). Further analyses were conducted with highly expressed genes with ≥ 10 TPM on a single transcript or multiple isoform basis. High fold upregulated and downregulated expression were considered if the fold changes were ≥ 10; FDR p≤0.05. To indicate differential expression on a single transcript level, we use the conventional term “differentially expressed genes” (DEGs). The differentially expressed transcript isoforms are referred to as DETI. In addition, transcript isoform switching was considered when a transcript was expressed with ≥ 10 TPM in either mGCs or cGCs, and the other group expressed with <0.05 TPM, so that the fold changes were ≥ 200; FDR p≤0.05.

### 2.7. Regulators of transcript isoforms

We carefully analyzed four different lists: (a) 651 mouse epigenetic regulators (41) (b) 2777 mouse Transcription factors (42) (c) 2792 mouse RNA-binding proteins (RBPs) (43, 44) (d) 85 mouse spliceosome-associated proteins (SPs) (45, 46),(e) 99 mouse transcription terminators and polyadenylation factors, which were compiled from (47). New tracks containing only these four groups were generated from each RNA-Seq data file that included expression values of multiple isoforms. These tracks were then used in subsequent analyses to identify the DETI (40).

### 2.8. Statistical analysis

In the CLC Genomics Workbench, the ‘Differential Expression for RNA-Seq’ tool performs multivariate statistics on a set of expression tracks based on a negative binomial generalized linear model (GLM). The final GLM fit and dispersion estimate determine the overall likelihood of the model given the data, as well as the uncertainty of each fitted coefficient. Two statistical tests—the Wald and the Likelihood Ratio tests—use one of these values. The across-groups (ANOVA-like) comparison employs the Likelihood Ratio test. RT-qPCR data were analysed using IBM SPSS

## 3. RESULTS

### 3.1. Mural and cumulus granulosa cells

Before analyzing the RNA-seq data, we assessed the reproducibility of the experimental model. Two hours after administering hCG to eCG-primed mice, we observed synchronized development of ovarian follicles to the antral stage, making it easier to isolate mGCs and cGCs for RNA extraction and transcriptome analysis. Within these antral follicles, two distinct populations of GCs were evident: mGCs forming layers lining the antral cavity and cGCs organized around the oocytes (**Fig. 1A-C**). Analyses of mGC and cGC transcriptomes revealed a group-specific correlation between the samples and transcript expression on either a single transcript or multiple isoform basis (**Fig. 1D-G**). RNA-Seq data were screened for GC-specific core gene expression. We observed that the overall expression pattern correlated with the previous reports on mGC or cGC-specific gene expression (**Fig. 1H** and **Supplementary Table S2**). Transcription factors (TFs) that have been used for reprogramming of human fibroblast or iPS cells to GC-like cells were expressed at high levels in mGCs and downregulated in cGCs, suggesting that cGCs are more differentiated compared to mGCs (**Fig. 1I-M**). We also performed RT-qPCR analyses of characteristic mGC- or cGC-specific gene expression, which indicates the purity of the isolated cells. These results demonstrate the reproducibility of the experimental model for studying mural and cumulus cells (**Fig. 1N-R**).

**Figure 1.**
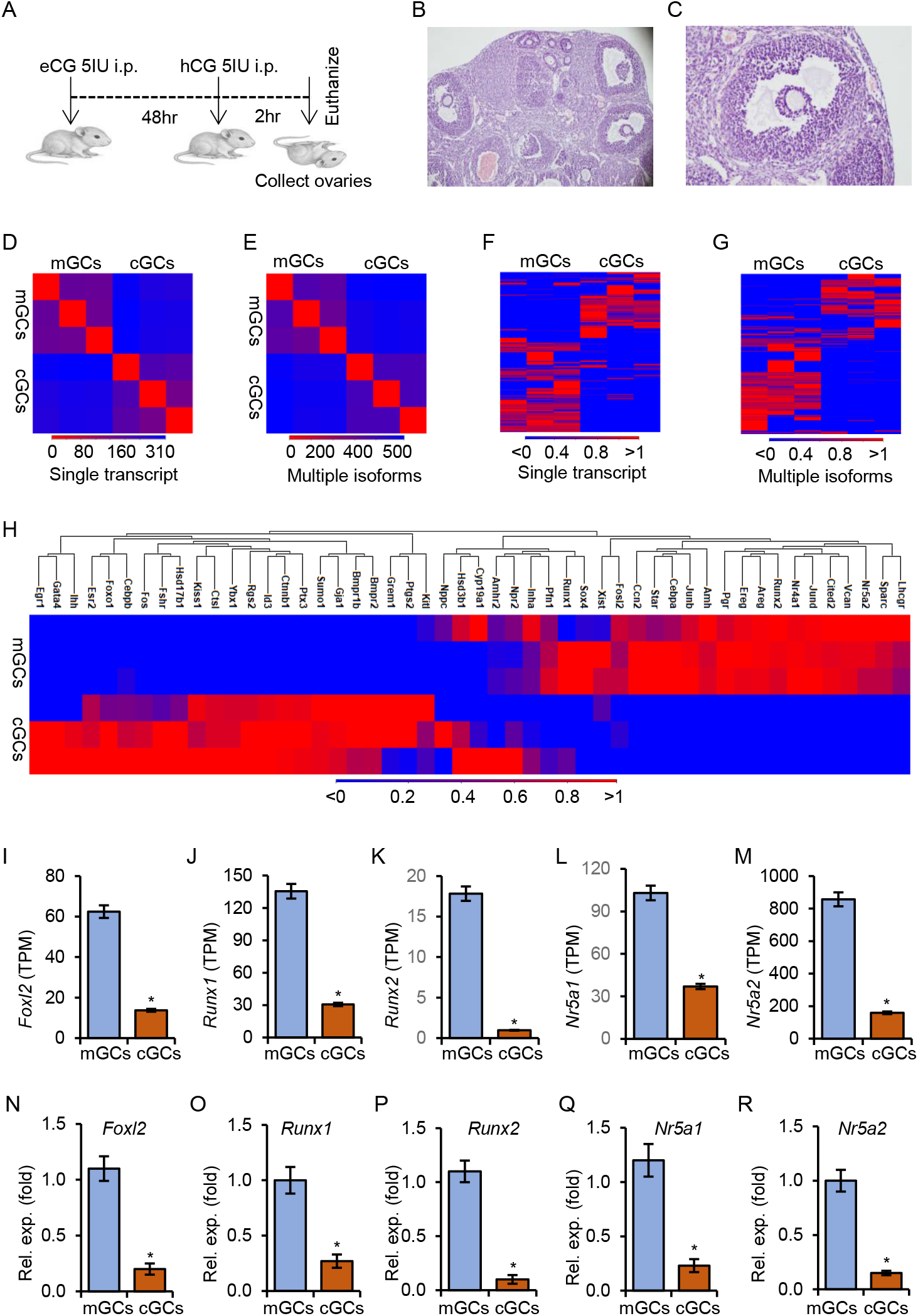
Mural and cumulus cell isolation, RNA sequencing, and data validation. A). Three-week-old wildtype C57BL/6 mice were intraperitoneally (i.p.) injected with 5IU of eCG. 48h after eCG injection, mice were injected with 5IU of hCG and euthanized 2 h after hCG injection to collect the ovaries and isolate the mural and cumulus granulosa cells. B, C). Ovarian histology showing the development of antral follicles. The mural and cumulus granulosa (mGC and cGC) specific transcriptomes were detected by RNA-sequencing, and the data were analyzed on either a single transcript or a multiple transcript isoform basis. Heat maps showing transcript expression at the sample level (D, E) and the transcript level (F, G). The single transcript-based expression pattern in mGC and cGC correlated with the previous reports (H). The transcription factors (TFs) used to reprogram human fibroblasts or iPS cells into GC-like cells were expressed at high levels in mGCs and downregulated in cGCs (I-M). The transcriptomic data also correlated well with RT-qPCR analyses (N-R).

### 3.2. Differentially expressed genes and mRNA isoforms

On a single transcript basis, 15539 genes were expressed in mGCs or cGCs. Of these transcripts, 68% (10,612/15539) were differentially expressed between mGCs and cGCs (absolute fold change, aFC≥2; FDR p≤ 0.05) (**Fig. 2A, 2B**). Of these, 21% (1816/8815) of the abundant transcripts (≥10TPM) showed a high-fold differential expression (aFC≥10; FDR p≤ 0.05) (**Fig. 2C, 2D)**. The genes such as *Pgr, Runx2, Nr4a1, Dot1l, Satb2, Elf4, Amh, Lhcgr, Areg, Ihh, Hhip, Gsk3a*, and *Prmt1* werer downregulated in cGCs. On the other hand, genes like *Id3, Yy2, Rbm7, Med6, Sumo1, Sumo2, Aurka, Hprt1, Rheb, Fkbp3, Fkbp7, Med10, Kiss1*, and *Cdkn3* showed upregulation in cGCs (**Supplementary Tables S3 and S4**). On a multiple transcript isoform basis, 39118 isoforms were expressed (≥1TPM) in mGCs or cGCs. Overall, 70% (27386/39118) of the transcript isoforms were differentially expressed (aFC≥2; FDR p≤ 0.05) (**Fig. 2E, 2F**). Among the abundant isoforms (≥10TPM), over 43% (5542/12854) showed a high-fold differential expression (aFC≥10; FDR p≤ 0.05) (**Fig. 2G, 2H**, and **Supplementary Tables S5 and S6**). Given that a large number of transcripts were differentially expressed between mGCs and cGCs, we focused on those that were abundantly expressed and exhibited a high level of differential expression.

**Figure 2.**
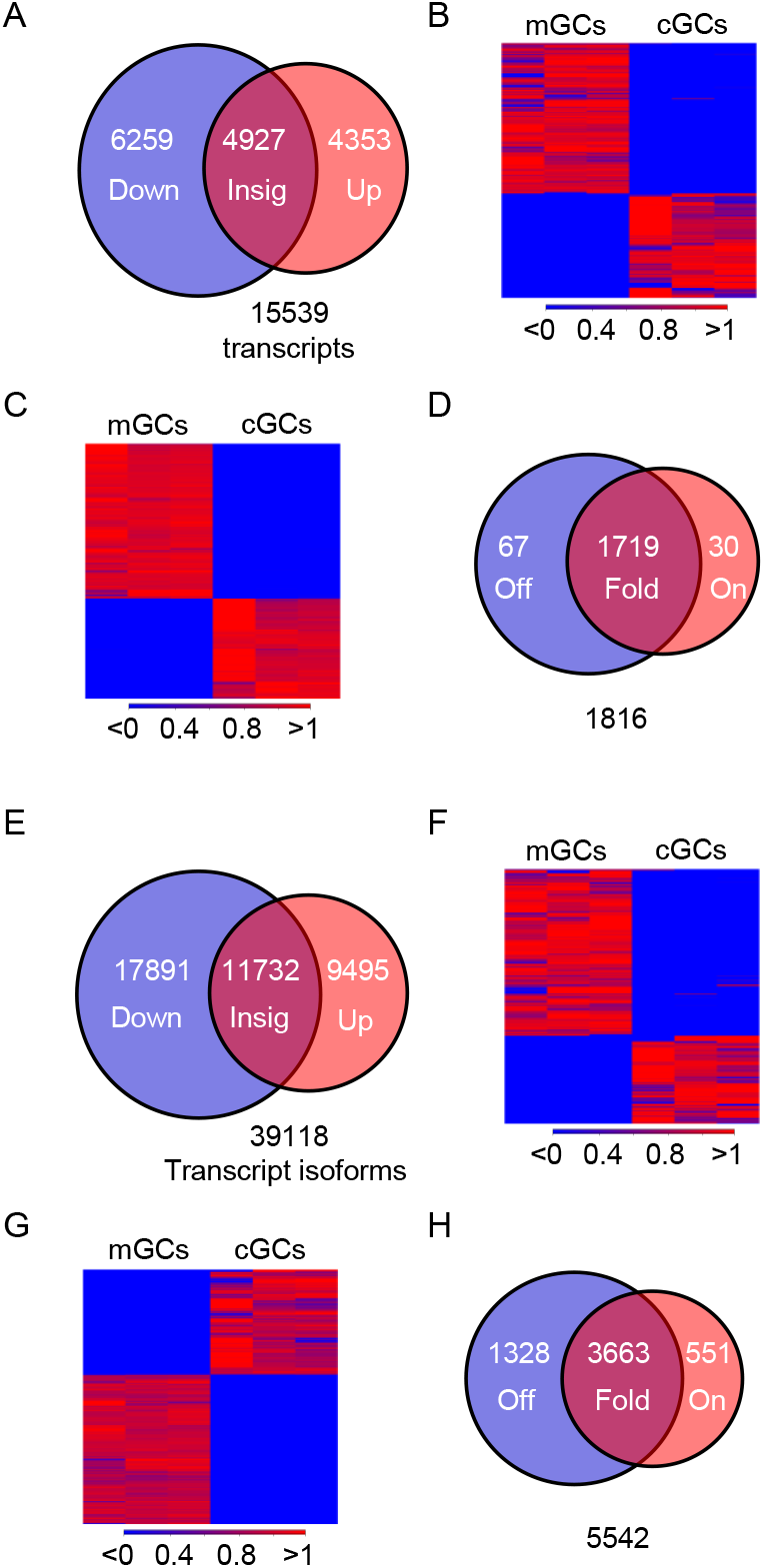
Differential expression of genes. On a single transcript basis, 15539 genes were expressed with ≥1TPM in mGCs or cGCs. Of these transcripts, 68% (10,612/15539) were differentially expressed in cGCs (aFC≥2; FDR p≤ 0.05) (A, B). Among the abundant transcripts with ≥10 TPM, 21% (1816/8815) showed a high-fold differential expression (aFC≥10; FDR p≤ 0.05). Remarkably, 67 transcripts were highly repressed (Off) and 30 were induced (On) in cGCs (C, D). On a multiple transcript isoform basis, 39118 isoforms were expressed (≥ 1TPM) in mGCs or cGCs. Overall, 70% of the isoforms (27386/39118) were differentially expressed (E, F), and among the abundant isoforms (≥10 TPM), over 43% (5542/12854) showed a high-fold differential expression (aFC≥10; FDR p≤ 0.05) (G). Of these 5542 differentially expressed transcript isoforms, 1328 were highly repressed and 551 were induced in cGCs (H).

### 3.3 Coding and noncoding transcripts

We have identified both DEGs and DETIs (**Fig. 2** and **Supplementary Tables S3-S6**). Among the 714 upregulated DEGs (TPM ≥ 10, fold-changes ≥ 10) in cGCs, 639 (89.5%) were protein-coding, and 62 (8.7%) were lncRNA. In contrast, among the 1102 downregulated DEGs, 1003 (91.07%) were protein-coding, and 93 (8.4%) were lncRNA. We also analyzed the biotype of DETIs. Among the 2280 upregulated DETIs, 1718 (75.4%) were protein-coding, 94 (4.1%) were lncRNA, 193 (8.5%) were destined to NMD, and 270 (11.8%) were retaining introns. Among the 3262 downregulated DETIs, 2606 (79.9%) were protein-coding, 134 (4.2%) were lncRNA, 116 (3.5%) were destined to NMD, and 404 (12.4%) were retaining introns.

### 3.4. Correlation between differentially expressed genes and transcripts

*The pattern of gene expression observed on a single transcript basis did not closely match their* respective multiple transcript isoforms. Among the 1816 DEGs with abundance and high fold changes, 714 were upregulated and 1102 were downregulated in cGCs. The 714 upregulated genes expressed 2,067 transcript isoforms across both cell types, with 22% (573/2,067) being either downregulated or showing insignificant changes in cGCs (**Fig. 3A, 3B**). Similarly, 1102 downregulated genes expressed 3,202 transcript isoforms, of which 4% (127/3202) were upregulated or showed no significant changes (**Fig. 3C, 3D**). Remarkably, the 4927 genes that showed no significant changes in expression at the single transcript level contained over 50% of their transcript isoforms that were differentially expressed (7103 ≥2 fold, ≥1 TPM, and 1218 ≥10 fold, ≥10 TPM; FDR p≤ 0.05) (**Fig. 3E, 3F**). These included transcripts of *Chad1, Esr2, Ezh2, Fos, Gata4, Kdm5a/5b, Myc*, and *Ybx1*. These findings suggest that the actual differential expression of transcripts may be missed if RNA-Seq data is only analyzed considering that a gene expresses a single transcript. Therefore, in the following sections, we focus on DETIs.

**Figure 3.**
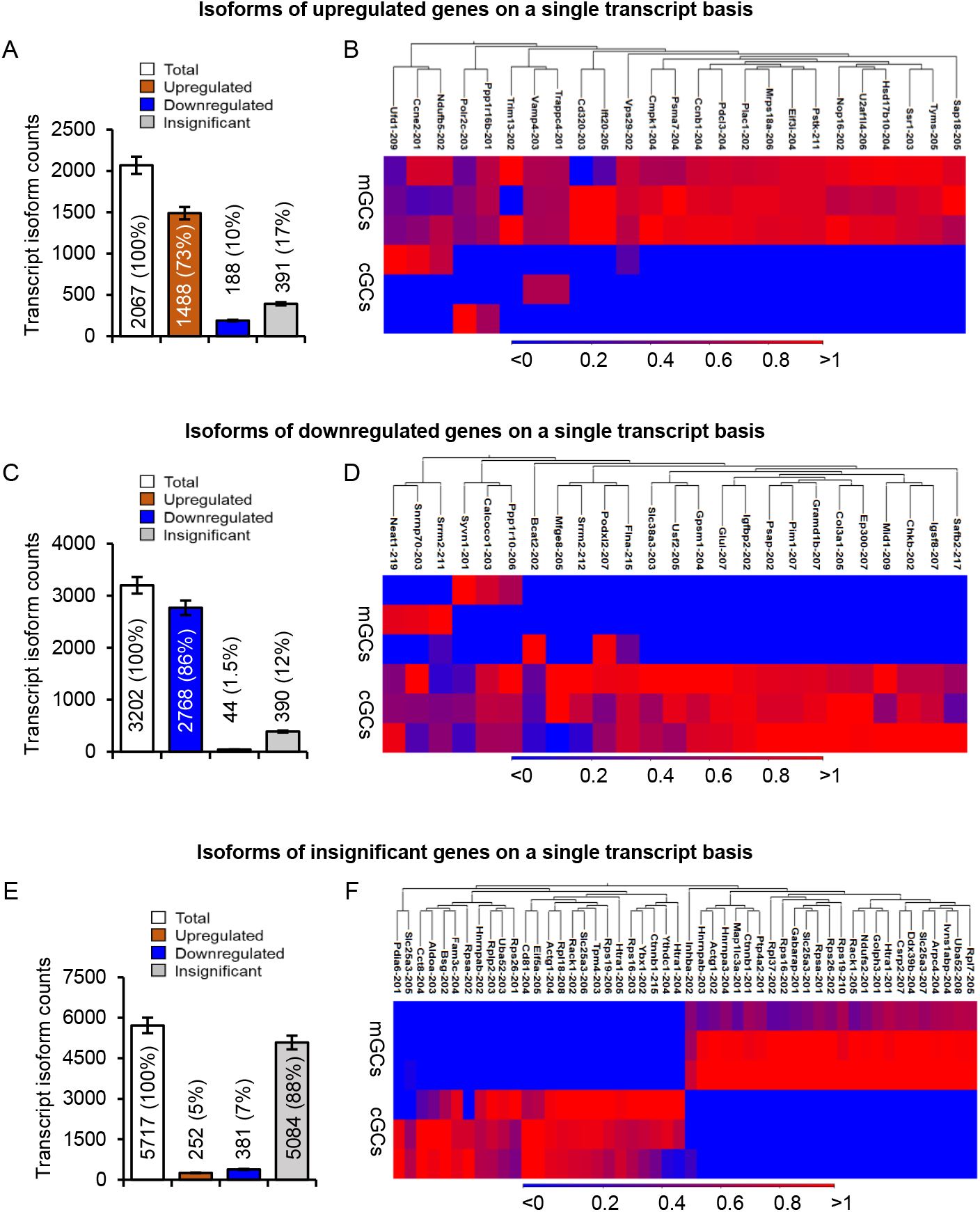
Isoforms of differentially expressed genes on a single transcript basis. Genes that did not show any differential expression (insignificant) on a single transcript basis expressed transcript isoforms that were significantly upregulated or downregulated (A, B). Similarly, some genes found upregulated on a single transcript basis expressed transcript isoforms that were either downregulated or insignificant (C, D). Genes found downregulated on a single transcript basis also expressed transcript isoforms that were either upregulated or insignificant (E, F). The bar graphs show the distribution of the transcript isoform counts (A, C, E), and the heatmaps show 50 differentially expressed isoforms in the insignificant group (B), 25 downregulated transcript isoforms in the upregulated group (D), and 25 upregulated isoforms in the downregulated group (F).

### 3.5. Unfolding of the transcript switching

Further analysis revealed that a group of transcript isoforms was switched off or switched on in cGCs (≥ 10 TPM in one group and < 0.05 TPM in the other) (**Fig. 4A** and **Supplementary Table S7)**. Some isoforms of *Adar, Cebpa, Dnmt3a, Foxo4, Pgr, Rest, Runx1, Satb2, Sirt1, Sirt2*, and *Tead1* were switched off, while isoforms of *Brd7, Crem, Chd1, Med21, Med27, Nfkbia, Rbm39, Rbmx, Suv39h2, Tcf12, Xist*, and *Ybx3* were switched on in cGCs. Additionally, 57 genes, including *Csde1, Dab2, Ezh2, Gata4, Gnas, Gtf2i, Macf1, Klf10, Setdb1*, and *Sp3*, had one isoform turned off at the same time as another isoform was turned on in cGCs (**Fig. 4B** and **Supplementary Table S8)**.

**Figure 4.**
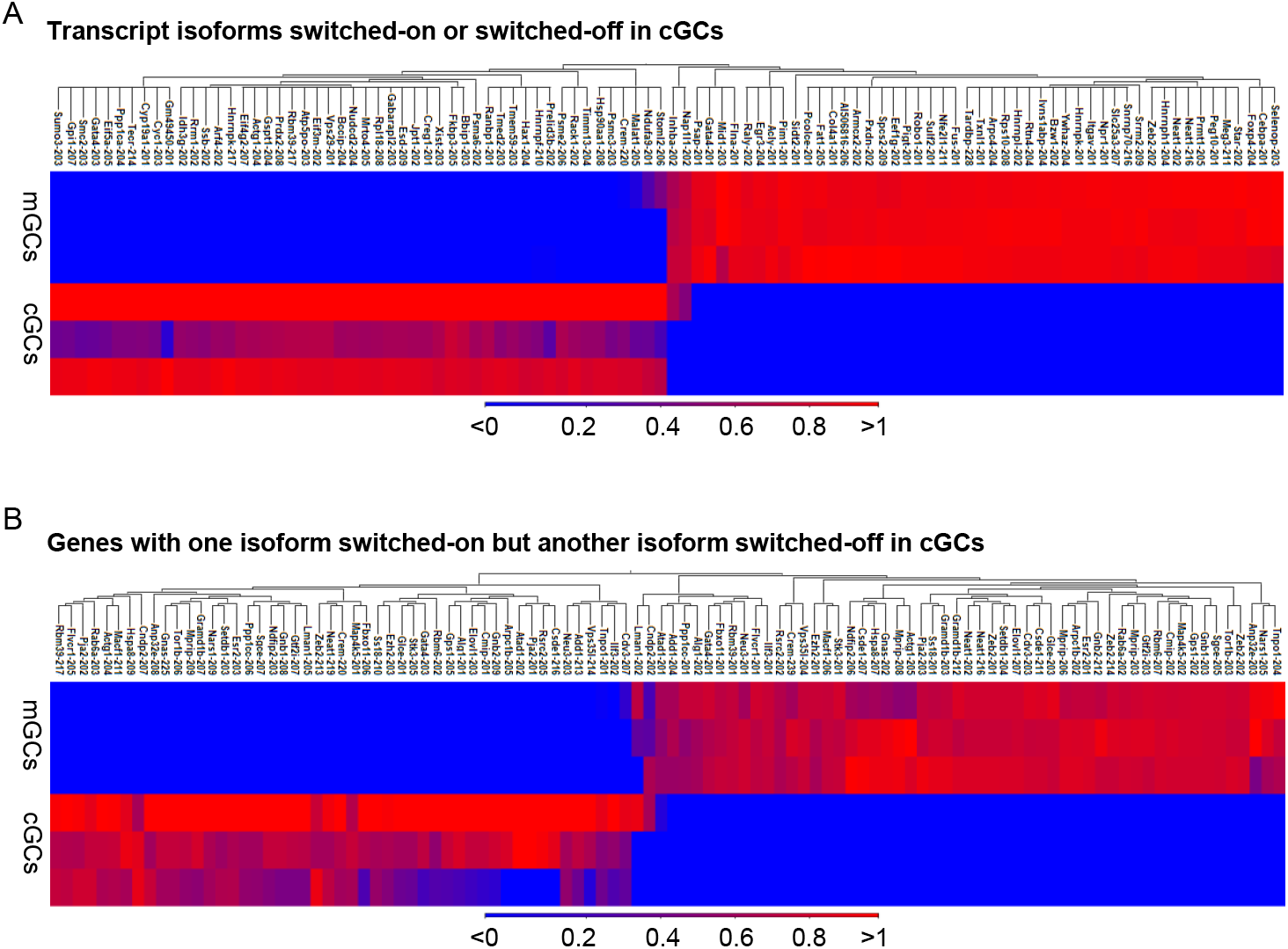
Transcript switching in mural and cumulus cells. Among the highly differentially expressed transcripts in mGCs and cGCs, one-third showed transcript switching (≥10 TPM in mGC or cGC but ≥200-fold lower in the other). The top 50 switched-on and top 50 switched-off transcripts are shown in the upper panel (A). The low panel included the isoforms of 57 genes, which contained one isoform switched on and another transcript switched off in cGCs (B).

### 3.6. Potential regulators of transcript switching

To understand the mechanism behind alternative TSSs, we examined the differential expression of epigenetic regulators (**Fig. 5A** and **Supplementary Table S9)** and TFs (**Fig. 5B** and **Supplementary Table S10)**. We identified 303 DETIs of epigenetic regulators (TPM ≥ 10, aFC ≥ 10), of which 135 showed transcript switching. We also observed 1201 DETIs of TFs (TPM ≥ 10, aFC ≥ 10), of which 442 showed isoform switching in cGCs. To understand the mechanisms behind alternative splicing, we analyzed the differential expression of RBPs, including SPs and RNA modifiers (**Fig. 5C** and **Supplementary Table S11)**. We identified 1799 DETIs of RBPs in CGCs. (TPM ≥ 10, aFC ≥ 10). Among the DETIs, significant isoform switching was observed in 607 transcripts. To understand the mechanisms underlying alternative polyadenylation sites, we examined the differential expression of transcription terminators (TTs) and polyadenylation factors (PAFs) (**Fig. 5D** and **Supplementary Table S12)**. We identified 39 DETIs of TTs and PAFs (TPM ≥ 10, aFC ≥ 10), of which 9 transcripts were switched off in cGCs.

**Figure 5.**
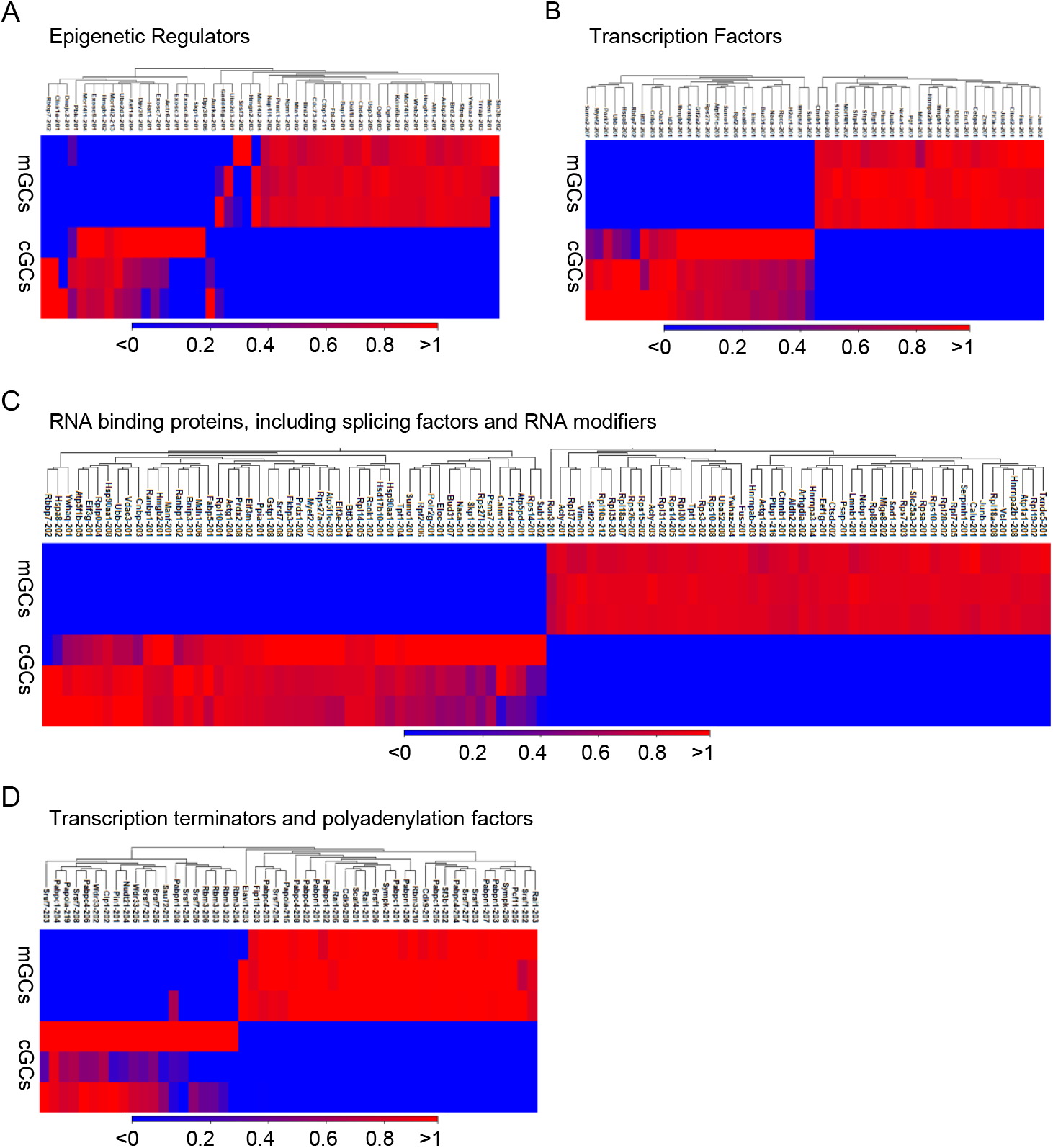
Differentially expressed regulators of transcript switching in cumulus granulosa cells. The upper panel displays the top 50 differentially expressed epigenetic regulators (A) and transcription factors (B). The middle panel shows the top 100 differentially expressed RNA-binding proteins, including splicing factors and RNA modifiers (C), and the lower panel shows the top 50 differentially expressed transcription terminators and polyadenylation factors (D).

## 4. DISCUSSION

Initially, we analyzed the RNA-seq data using the traditional ‘one gene-one transcript’ model; then, we examined transcript isoforms, considering that ‘one gene expresses various isoforms’ depending on cellular differentiation. When comparing the transcriptomes of mGCs and cGCs, the first question was which cell type is closer to the original GCs before gonadotropin-induced differentiation. Several recent studies have identified GC-specific core TFs that can reprogram ES cells or fibroblasts into granulosa-like cells (48, 49). The expression of these TFs was high in mGCs and significantly downregulated in cGCs (**Fig.1I-1R**). Therefore, we used the mGCs as controls to compare the changes in gene expression in cGCs. However, we acknowledge that RNA-seq data from GCs before gonadotropin stimulation would serve as a more suitable control for both mGCs and cGCs.

The RNA-seq data reveal the transcriptomic landscapes of mGCs and cGCs. At both the single transcript and multiple isoform levels, the number of transcripts expressed with similar TPM values was significantly lower in cGCs, indicating that overall transcription is more active in mGCs. Although a larger proportion (∼70%) of transcripts were downregulated in cGCs, this was not due to technical artifacts or RNA quality issues, as we also identified a substantial number (∼30%) of transcripts that were markedly upregulated in cGCs. These findings suggest that gonadotropin-induced differentiation may suppress overall transcription in cGCs while selectively upregulating specific genes. A previous study has shown that oocyte-derived GDF9 represses *Foxl2* expression in cGCs (50). We also observed that many TFs were downregulated in cGCs; however, further research is needed to uncover the underlying mechanisms.

While most transcripts are protein-coding, with or without a defined coding sequence, some include transcripts that undergo nonsense-mediated decay, retain introns, or are long non-coding RNAs. Some genes produce both coding and noncoding RNAs, yet they are classified as protein-coding in single transcript-based analyses. Without analyzing transcript isoforms, these noncoding RNAs would remain undetected. We identified new transcripts of both coding and noncoding RNAs, especially lncRNAs, that regulate gonadotropin-induced mural and cumulus cell differentiation. We acknowledge that further studies are needed to clarify the role of regulatory noncoding RNAs in GC differentiation.

Understanding the difference between single-transcript-based and multiple transcript-isoform-based differential expression is crucial, given the variety of transcript isoforms expressed during cell differentiation. When analyzing the expression of a single transcript, many genes appear to be strongly upregulated or downregulated (Fig. 2A-2D). Both single-transcript and multiple-isoform analyses showed that about 70% of transcripts are differentially expressed between mGCs and cGCs. However, their expression patterns don’t always match their transcript isoforms; for example, the genes that are upregulated may express transcript isoforms that are downregulated in cGCs, and vice versa (Fig. 3A-3D). Moreover, genes that show no significant changes in single-transcript analyses still exhibit a large number of DETIs (Fig. 3E-3F). When a gene has one transcript isoform that is significantly upregulated and another that is downregulated in cGCs, the overall gene expression in single transcript-based analyses might not appear significantly different. These results suggest that single transcript-based DEGs often do not reflect actual changes across transcript isoforms. Even if a single-transcript analysis indicates no change, specific isoforms can still be strongly up- or downregulated.

Both mGCs and cGCs differ from GCs in their responses to gonadotropins, so it is expected that both cell types will share many genes. As expected, over 90% of transcripts are common between them, but their expression levels vary during cell differentiation. We identified about 70% of all transcripts that showed a 2-fold to 1,000-fold change in expression; some were highly expressed in mGCs, while others demonstrated that, although one isoform of a gene is upregulated in cGCs, another isoform of the same gene can be upregulated in mGCs. Additionally, a gene can produce one transcript isoform at a much higher level, while another isoform is expressed at a significantly lower level in the same cell type. This kind of differential expression remains unclear in single-transcript-based analyses.

Differential expression of transcript isoforms also involves transcript switching, where one isoform can be suppressed entirely or a new isoform can be significantly upregulated during cell differentiation. Several transcript switching events have been previously reported during stem cell differentiation and tumorigenesis (51, 52). In this study, we observed a remarkable switching of transcript isoforms between mGCs and cGCs (Fig. 4A, 4B). The isoforms are differentially expressed due to cell-type-specific changes in epigenetic and transcriptional regulators, RBPs, including SFs, TTs, and PAFs, which are associated with alternative TSS, alternative splicing, and alternative polyadenylation sites (Fig. 5A-5D). Our findings show that gonadotropin-induced transcript switching in GCs is essential for the differentiation of mural and cumulus granulosa cells (Fig. 6A, 6B), which might have gone unnoticed without mRNA isoform analysis.

**Figure 6.**
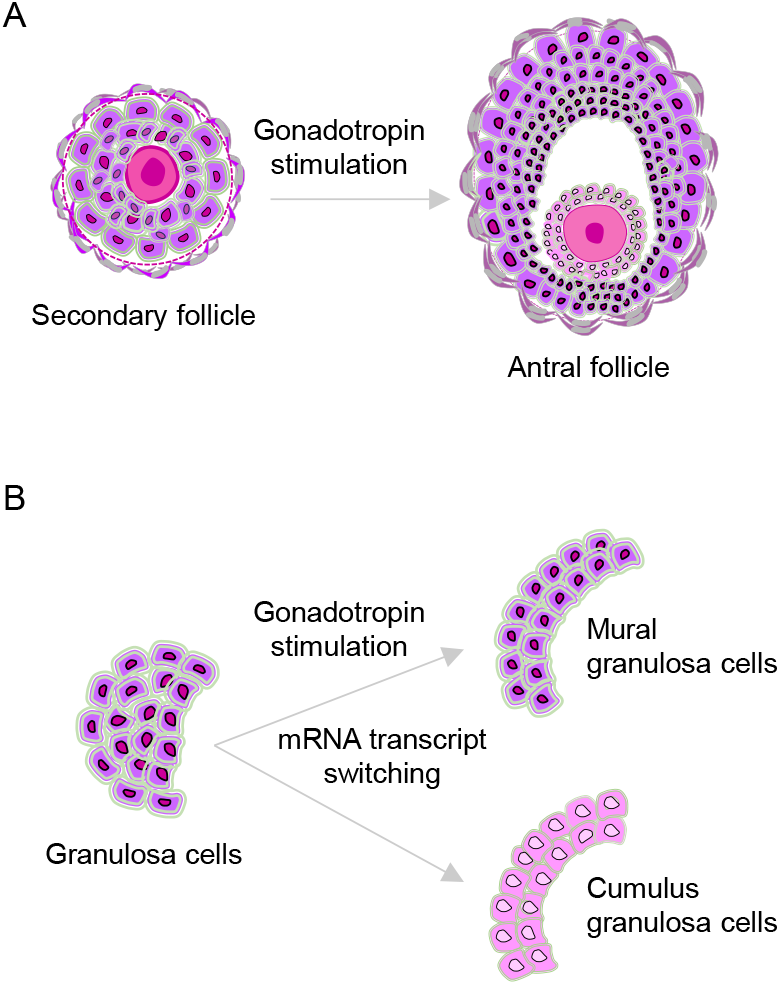
Mural and cumulus granulosa differentiation. Schematic presentation of gonadotropin-induced differentiation of granulosa cells in secondary follicles to mural and cumulus cells in antral follicles (A). Analyses of our results indicate that gonadotropin-induced transcript switching in granulosa cells plays a crucial role in mural and cumulus granulosa differentiation, which would remain undetected without the mRNA isoform analysis (B).

## Acknowledgements

We acknowledge the Department of Pathology and Laboratory Medicine at KUMC for supporting M.A.K.R. and P.E.F.

