## Supplementary Tables for "mRNA isoform switching plays a crucial role in mural cumulus differentiation"

Table S1. Primers used for conventional RT-PCR and RT-qPCR analyses

| Sl. | Gene | Ref. Seqs. | Forward primer | Reverse primer | PCR product |
|---|---|---|---|---|---|
| 1 | *Foxl2* | NM_012020.3 | mFoxl2-13F:<br>GCCAGAGGCTGACTTCCA | mFoxl2-353R:<br>GGTTTCTCCGGTGTTGTCC | 341 bp |
| 2 | *Runx1* | NM_009821.3<br>NM_001111023.2<br>NM_001111022.2<br>NM_001111021.2 | mRunx1-1947F:<br>TGAAGAACCAGGTAGCGAGA | mRunx1-2195R:<br>GGTGGACAGAGGAAGAGGTG | 249 bp |
| 3 | *Runx2* | NR_073425.2<br>NR_073392.2<br>NM_001271621.2<br>NM_001271630.2<br>NM_001145920.3<br>NM_009820.6<br>NM_001271627.2 | mRunx2-990F:<br>TACCTGCCATCACTGACGTG | mRunx2-1145R:<br>ATGAAATGCTTGGGAACTGC | 156 bp |
| 4 | *Nr5a1* | NM_001316687.1<br>NM_139051.3 | mNr5a1-1178F:<br>ATCTACCGCCAGGTCCAGTA | mNr5a1-1412R:<br>GGCTGTGGTTGTTCAGGAAT | 235 bp |
| 5 | *Nr5a2* | XM_006529621.3<br>XM_006529622.4<br>NM_030676.3<br>NM_001159769.2 | mNr5a2-1285F:<br>TTGAGTGGGCCAGGAGTAGT | mNr5a2-1485F:<br>ACGCGACTTCTGTGTGTGAG | 201 bp |

Sl., serial number; Ref. Seqs., reference sequences (NCBI)

Table S2. Expression of 50 known genes in mural (mGC) and cumulus (cGC) cells

| Name | ENSEMBL | Chrom | Exp in mGC (TPM) | Exp in cGC (TPM) | Fold change | FDR p-value | Biotype |
|---|---|---|---|---|---|---|---|
| *Amh* | ENSMUSG00000035262 | 10 | 407.17 | 1.15 | -353.49 | 0.00 | Protein coding |
| *Jund* | ENSMUSG00000071076 | 8 | 532.75 | 2.55 | -208.82 | 0.00 | Protein coding |
| *Nr4a1* | ENSMUSG00000023034 | 15 | 1336.17 | 16.00 | -83.50 | 0.00 | Protein coding |
| *Pgr* | ENSMUSG00000031870 | 9 | 332.00 | 5.42 | -61.20 | 0.00 | Protein coding |
| *Ereg* | ENSMUSG00000029377 | 5 | 2960.88 | 81.05 | -36.53 | 0.00 | Protein coding |
| *Junb* | ENSMUSG00000052837 | 8 | 1051.67 | 34.92 | -30.12 | 0.00 | Protein coding |
| *Ccn2* | ENSMUSG00000019997 | 10 | 850.96 | 32.17 | -26.45 | 0.00 | Protein coding |
| *Star* | ENSMUSG00000031574 | 8 | 2409.02 | 98.77 | -24.39 | 0.00 | Protein coding |
| *Vcan* | ENSMUSG00000021614 | 13 | 1216.58 | 59.64 | -20.40 | 0.00 | Protein coding |
| *Runx2* | ENSMUSG00000039153 | 17 | 17.83 | 0.97 | -18.33 | 0.00 | Protein coding |
| *Areg* | ENSMUSG00000029378 | 5 | 4895.02 | 272.55 | -17.96 | 0.00 | Protein coding |
| *Lhcgr* | ENSMUSG00000024107 | 17 | 452.50 | 25.38 | -17.83 | 0.00 | Protein coding |
| *Sparc* | ENSMUSG00000018593 | 11 | 427.14 | 29.50 | -14.48 | 0.00 | Protein coding |
| *Cited2* | ENSMUSG00000039910 | 10 | 1291.90 | 90.09 | -14.34 | 0.00 | Protein coding |
| *Cebpa* | ENSMUSG00000034957 | 7 | 528.22 | 38.98 | -13.55 | 0.00 | Protein coding |
| *Nr5a2* | ENSMUSG00000026398 | 1 | 857.22 | 159.04 | -5.39 | 0.00 | Protein coding |
| *Pfn1* | ENSMUSG00000018293 | 11 | 398.27 | 78.25 | -5.09 | 0.00 | Protein coding |
| *Sox4* | ENSMUSG00000076431 | 13 | 526.48 | 114.20 | -4.61 | 0.00 | Protein coding |
| *Xist* | ENSMUSG00000086503 | X | 184.61 | 41.58 | -4.44 | 0.00 | lncRNA |
| *Runx1* | ENSMUSG00000022952 | 16 | 135.43 | 30.64 | -4.42 | 0.00 | Protein coding |
| *Inha* | ENSMUSG00000032968 | 1 | 25533.19 | 5910.46 | -4.32 | 0.00 | Protein coding |
| *Fosl2* | ENSMUSG00000029135 | 5 | 374.44 | 109.17 | -3.43 | 0.00 | Protein coding |
| *Amhr2* | ENSMUSG00000023047 | 15 | 172.83 | 54.69 | -3.16 | 0.00 | Protein coding |
| *Npr2* | ENSMUSG00000028469 | 4 | 217.65 | 78.57 | -2.77 | 0.00 | Protein coding |

| | | | | | | | |
|---|---|---|---|---|---|---|---|
| *Nppc* | ENSMUSG00000026241 | 1 | 167.49 | 65.43 | -2.56 | 0.00 | Protein coding |
| *Cyp19a1* | ENSMUSG00000032274 | 9 | 604.27 | 270.97 | -2.23 | 0.00 | Protein coding |
| *Egr1* | ENSMUSG00000038418 | 18 | 1411.17 | 698.60 | -2.02 | 0.00 | Protein coding |
| *Kitl* | ENSMUSG00000019966 | 10 | 401.82 | 202.94 | -1.98 | 0.00 | Protein coding |
| *Hsd3b1* | ENSMUSG00000027871 | 3 | 7033.18 | 3951.22 | -1.78 | 0.01 | Protein coding |
| *Ihh* | ENSMUSG00000006538 | 1 | 175.43 | 119.34 | -1.47 | 0.06 | Protein coding |
| *Esr2* | ENSMUSG00000021055 | 12 | 141.36 | 96.82 | -1.46 | 0.03 | Protein coding |
| *Ptgs2* | ENSMUSG00000032487 | 1 | 915.67 | 789.37 | -1.16 | 0.56 | Protein coding |
| *Ctnnb1* | ENSMUSG00000006932 | 9 | 415.73 | 374.53 | -1.11 | 0.58 | Protein coding |
| *Cebpb* | ENSMUSG00000056501 | 2 | 138.80 | 133.46 | -1.04 | 0.87 | Protein coding |
| *Ybx1* | ENSMUSG00000028639 | 4 | 1917.17 | 1936.34 | 1.01 | 0.95 | Protein coding |
| *Gata4* | ENSMUSG00000021944 | 14 | 121.32 | 138.31 | 1.14 | 0.56 | Protein coding |
| *Foxo1* | ENSMUSG00000044167 | 3 | 142.17 | 172.03 | 1.21 | 0.34 | Protein coding |
| *Bmpr1b* | ENSMUSG00000052430 | 3 | 90.03 | 158.45 | 1.76 | 0.01 | Protein coding |
| *Fos* | ENSMUSG00000021250 | 12 | 185.12 | 351.73 | 1.90 | 0.00 | Protein coding |
| *Bmpr2* | ENSMUSG00000067336 | 1 | 158.34 | 311.93 | 1.97 | 0.00 | Protein coding |
| *Gja1* | ENSMUSG00000050953 | 10 | 937.22 | 2043.15 | 2.18 | 0.00 | Protein coding |
| *Grem1* | ENSMUSG00000074934 | 2 | 61.45 | 135.20 | 2.20 | 0.00 | Protein coding |
| *Fshr* | ENSMUSG00000032937 | 17 | 66.19 | 148.92 | 2.25 | 0.00 | Protein coding |
| *Ptx3* | ENSMUSG00000027832 | 3 | 56.03 | 206.74 | 3.69 | 0.00 | Protein coding |
| *Rgs2* | ENSMUSG00000026360 | 1 | 743.89 | 2878.84 | 3.87 | 0.00 | Protein coding |
| *Hsd17b1* | ENSMUSG00000019301 | 11 | 349.93 | 1676.16 | 4.79 | 0.00 | Protein coding |
| *Ctsl* | ENSMUSG00000021477 | 13 | 721.81 | 3637.90 | 5.04 | 0.00 | Protein coding |
| *Id3* | ENSMUSG00000007872 | 4 | 218.68 | 2746.57 | 12.56 | 0.00 | Protein coding |
| *Sumo1* | ENSMUSG00000026021 | 1 | 37.92 | 2094.61 | 55.24 | 0.00 | Protein coding |
| *Kiss1* | ENSMUSG00000116158 | 1 | 0.02 | 11.12 | 680.36 | 0.00 | Protein coding |

Chrom, chromosome; Exp, expression; TPM, transcript per million; FDR, false discovery rate; NMD, nonsense mediated decay

Table S3. Top 50 upregulated genes in cumulus granulosa cells

| Name | ENSEMBL | Chrom | Exp in mGC (TPM) | Exp in cGC (TPM) | Fold change | FDR p-value | Biotype |
|---|---|---|---|---|---|---|---|
| *Rbm7* | ENSMUSG00000042396 | 9 | 13.39 | 134.17 | 10.02 | 0.00 | Protein coding |
| *Id3* | ENSMUSG00000007872 | 4 | 82.23 | 970.33 | 11.80 | 0.00 | Protein coding |
| *Rfc4* | ENSMUSG00000022881 | 16 | 218.68 | 2746.57 | 12.56 | 0.00 | Protein coding |
| *Med6* | ENSMUSG00000002679 | 12 | 11.51 | 162.13 | 14.09 | 0.00 | Protein coding |
| *Hat1* | ENSMUSG00000027018 | 2 | 10.10 | 155.73 | 15.42 | 0.00 | Protein coding |
| *Rpa3* | ENSMUSG00000012483 | 6 | 12.88 | 206.87 | 16.06 | 0.00 | Protein coding |
| *Med10* | ENSMUSG00000021598 | 13 | 18.42 | 300.83 | 16.33 | 0.00 | Protein coding |
| *Fkbp7* | ENSMUSG00000002732 | 2 | 7.27 | 158.94 | 21.86 | 0.00 | Protein coding |
| *Kiss1* | ENSMUSG00000116158 | 1 | 0.24 | 29.36 | 122.02 | 0.00 | Protein coding |
| *Psmg1* | ENSMUSG00000022913 | 16 | 0.12 | 15.95 | 132.04 | 0.00 | Protein coding |
| *Aurka* | ENSMUSG00000027496 | 2 | 0.23 | 30.71 | 133.56 | 0.00 | Protein coding |
| *Sumo2* | ENSMUSG00000020738 | 11 | 2.02 | 306.67 | 151.95 | 0.00 | Protein coding |
| *Ppa1* | ENSMUSG00000020089 | 10 | 4.12 | 741.40 | 180.07 | 0.00 | Protein coding |
| *Dut* | ENSMUSG00000027203 | 2 | 0.61 | 111.25 | 181.87 | 0.00 | Protein coding |
| *Eloc* | ENSMUSG00000079658 | 1 | 1.53 | 330.89 | 215.66 | 0.00 | Protein coding |
| *Dpy30* | ENSMUSG00000024067 | 17 | 1.21 | 281.55 | 233.10 | 0.00 | Protein coding |
| *Rpl9-ps6* | ENSMUSG00000062456 | 19 | 0.33 | 81.13 | 245.00 | 0.00 | Protein coding |
| *Erhrt-ps* | ENSMUSG00000063412 | 8 | 0.40 | 149.95 | 375.70 | 0.00 | Protein coding |
| *Lamtor5* | ENSMUSG00000087260 | 3 | 1.17 | 466.14 | 398.40 | 0.00 | Protein coding |
| *Bccip* | ENSMUSG00000030983 | 7 | 0.47 | 188.63 | 398.52 | 0.00 | Protein coding |
| *Psma1* | ENSMUSG00000030751 | 7 | 4.38 | 1841.89 | 420.30 | 0.00 | Protein coding |
| *Eif3m* | ENSMUSG00000027170 | 2 | 1.95 | 954.27 | 488.69 | 0.00 | Protein coding |
| *Rpf2* | ENSMUSG00000038510 | 10 | 0.02 | 12.48 | 514.42 | 0.00 | Protein coding |
| *Ndufs4* | ENSMUSG00000021764 | 13 | 0.96 | 510.26 | 529.74 | 0.00 | Protein coding |
| *Myef2* | ENSMUSG00000027201 | 2 | 0.86 | 515.96 | 602.06 | 0.00 | Protein coding |

| | | | | | | | |
|---|---|---|---|---|---|---|---|
| *Bud31* | ENSMUSG00000038722 | 5 | 2.74 | 1861.27 | 680.36 | 0.00 | Protein coding |
| *Polr2g* | ENSMUSG00000071662 | 19 | 0.23 | 155.77 | 682.66 | 0.00 | Protein coding |
| *Psmd14* | ENSMUSG00000026914 | 2 | 1.18 | 900.79 | 760.93 | 0.00 | Protein coding |
| *Tnni3* | ENSMUSG00000035458 | 7 | 0.36 | 331.21 | 923.46 | 0.00 | Protein coding |
| *Immp1l* | ENSMUSG00000042670 | 2 | 0.78 | 743.20 | 948.40 | 0.00 | Protein coding |
| *Cetn3* | ENSMUSG00000021537 | 13 | 0.97 | 1297.49 | 1337.26 | 0.00 | Protein coding |
| *Crabp2* | ENSMUSG00000004885 | 3 | 0.15 | 218.95 | 1485.95 | 0.00 | Protein coding |
| *Cacybp* | ENSMUSG00000014226 | 1 | 0.27 | 461.43 | 1704.99 | 0.00 | Protein coding |
| *Gstp1* | ENSMUSG00000060803 | 19 | 0.27 | 474.36 | 1775.30 | 0.00 | Protein coding |
| *Bnip3* | ENSMUSG00000078566 | 7 | 0.54 | 971.94 | 1807.80 | 0.00 | Protein coding |
| *Fam162a* | ENSMUSG00000003955 | 16 | 0.16 | 295.90 | 1861.29 | 0.00 | Protein coding |
| *Rps27rt* | ENSMUSG00000050621 | 9 | 0.30 | 678.48 | 2247.12 | 0.00 | Protein coding |
| *Ubb-ps_1* | ENSMUSG00000121785 | 14 | 0.30 | 712.06 | 2334.89 | 0.00 | lncRNA |
| *Hmgb2* | ENSMUSG00000054717 | 8 | 0.06 | 161.44 | 2516.88 | 0.00 | Protein coding |
| *Hprt1* | ENSMUSG00000025630 | X | 0.36 | 917.66 | 2531.28 | 0.00 | Protein coding |
| *Psmb2* | ENSMUSG00000028837 | 4 | 0.01 | 16.39 | 2804.81 | 0.00 | Protein coding |
| *Srsf7* | ENSMUSG00000024097 | 17 | 0.17 | 549.24 | 3199.47 | 0.00 | Protein coding |
| *Atp5f1c* | ENSMUSG00000025781 | 2 | 0.02 | 118.39 | 5861.76 | 0.00 | Protein coding |
| *Mdh1* | ENSMUSG00000020321 | 11 | 0.04 | 238.73 | 6348.43 | 0.00 | Protein coding |
| *Rps27l* | ENSMUSG00000036781 | 9 | 0.00 | 13.47 | 6712.16 | 0.00 | Protein coding |
| *Sumo1* | ENSMUSG00000026021 | 1 | 0.05 | 456.30 | 9148.76 | 0.00 | Protein coding |
| *Ranbp1* | ENSMUSG00000005732 | 16 | 0.01 | 62.08 | 10208.71 | 0.00 | Protein coding |
| *Prdx1* | ENSMUSG00000028691 | 4 | 0.01 | 94.32 | 14079.82 | 0.00 | Protein coding |
| *Rgcc* | ENSMUSG00000022018 | 14 | 0.00 | 22.31 | 24340.28 | 0.00 | Protein coding |
| *Ppia* | ENSMUSG00000071866 | 11 | 0.00 | 26.31 | 32220.91 | 0.00 | Protein coding |

Chrom, chromosome; Exp, expression; TPM, transcript per million; FDR, false discovery rate; NMD, nonsense mediated decay

Table S4. Top 50 downregulated genes in cumulus granulosa cells

| Name | ENSEMBL | Chrom | Exp in mGC (TPM) | Exp in cGC (TPM) | Fold change | FDR p-value | Biotype |
|---|---|---|---|---|---|---|---|
| *Edn2* | ENSMUSG00000028635 | 4 | 602.67 | 0.03 | -21544.91 | 0.00 | Protein coding |
| *Tnfsf11* | ENSMUSG00000022015 | 14 | 82.06 | 0.01 | -6000.30 | 0.00 | Protein coding |
| *Gsk3a* | ENSMUSG00000057177 | 7 | 82.06 | 0.01 | -6000.30 | 0.00 | Protein coding |
| *Proser1* | ENSMUSG00000049504 | 3 | 49.38 | 0.01 | -5404.89 | 0.00 | Protein coding |
| *Raver1* | ENSMUSG00000010205 | 9 | 108.51 | 0.05 | -2180.11 | 0.00 | Protein coding |
| *Ece1* | ENSMUSG00000057530 | 4 | 58.12 | 0.01 | -3890.85 | 0.00 | Protein coding |
| *Ppp1r10* | ENSMUSG00000039220 | 17 | 23.14 | 0.01 | -3393.76 | 0.00 | Protein coding |
| *Mier2* | ENSMUSG00000042570 | 10 | 48.07 | 0.02 | -2778.43 | 0.00 | Protein coding |
| *Fbrs* | ENSMUSG00000042423 | 7 | 20.96 | 0.01 | -2302.33 | 0.00 | Protein coding |
| *Uck2* | ENSMUSG00000026558 | 1 | 108.51 | 0.05 | -2180.11 | 0.00 | Protein coding |
| Akap17a | ENSMUSG00000121606 | X | 25.42 | 0.01 | -2105.92 | 0.00 | Protein coding |
| *Reck* | ENSMUSG00000028476 | 4 | 101.95 | 0.05 | -1915.50 | 0.00 | Protein coding |
| *Tmem115* | ENSMUSG00000010045 | 9 | 66.82 | 0.04 | -1684.34 | 0.00 | Protein coding |
| *Sf1* | ENSMUSG00000024949 | 19 | 23.59 | 0.01 | -1667.69 | 0.00 | Protein coding |
| *Ncs1* | ENSMUSG00000062661 | 2 | 369.88 | 0.23 | -1583.02 | 0.00 | Protein coding |
| *Eif3k* | ENSMUSG00000053565 | 7 | 55.14 | 0.04 | -1557.43 | 0.00 | Protein coding |
| *Etv5* | ENSMUSG00000013089 | 16 | 38.65 | 0.03 | -1391.99 | 0.00 | Protein coding |
| *Tbc1d2b* | ENSMUSG00000037410 | 9 | 17.27 | 0.01 | -1158.32 | 0.00 | Protein coding |
| *Imp3* | ENSMUSG00000032288 | 9 | 14.08 | 0.01 | -1155.70 | 0.00 | Protein coding |
| *Ptov1* | ENSMUSG00000038502 | 7 | 18.14 | 0.02 | -1153.55 | 0.00 | Protein coding |
| *Erbb2* | ENSMUSG00000062312 | 11 | 14.63 | 0.01 | -1130.45 | 0.00 | Protein coding |
| *Srbd1* | ENSMUSG00000024135 | 17 | 104.50 | 0.10 | -1097.85 | 0.00 | Protein coding |
| *Slc38a10* | ENSMUSG00000061306 | 11 | 75.22 | 0.07 | -1072.35 | 0.00 | Protein coding |
| *Cbx2* | ENSMUSG00000025577 | 11 | 15.80 | 0.01 | -1070.76 | 0.00 | Protein coding |
| *Pde7b* | ENSMUSG00000019990 | 10 | 11.22 | 0.01 | -1066.58 | 0.00 | Protein coding |
| *Arhgdia* | ENSMUSG00000025132 | 11 | 13.31 | 0.01 | -1004.81 | 0.00 | Protein coding |

| | | | | | | | |
|---|---|---|---|---|---|---|---|
| *Snn* | ENSMUSG00000037972 | 16 | 43.02 | 0.04 | -985.97 | 0.00 | Protein coding |
| *Zbtb39* | ENSMUSG00000044617 | 10 | 26.20 | 0.03 | -957.63 | 0.00 | Protein coding |
| *Ubald2* | ENSMUSG00000050628 | 11 | 10.24 | 0.01 | -910.39 | 0.00 | Protein coding |
| *Cyba* | ENSMUSG00000006519 | 8 | 34.95 | 0.04 | -881.28 | 0.00 | Protein coding |
| *Tead1* | ENSMUSG00000055320 | 7 | 60.20 | 0.07 | -880.95 | 0.00 | Protein coding |
| *Adgre5* | ENSMUSG00000002885 | 8 | 39.28 | 0.04 | -874.80 | 0.00 | Protein coding |
| *Gabarapl1* | ENSMUSG00000030161 | 6 | 39.99 | 0.05 | -859.36 | 0.00 | Protein coding |
| *Tmem94* | ENSMUSG00000020747 | 11 | 11.20 | 0.01 | -838.26 | 0.00 | Protein coding |
| *Eif3f* | ENSMUSG00000031029 | 7 | 50.51 | 0.06 | -802.06 | 0.00 | Protein coding |
| *Prnp* | ENSMUSG00000079037 | 2 | 10.50 | 0.01 | -771.44 | 0.00 | Protein coding |
| *Gorasp1* | ENSMUSG00000032513 | 9 | 32.15 | 0.04 | -748.73 | 0.00 | Protein coding |
| *Zc3h7b* | ENSMUSG00000022390 | 15 | 29.58 | 0.04 | -747.94 | 0.00 | Protein coding |
| *Baiap2* | ENSMUSG00000025372 | 11 | 60.82 | 0.08 | -738.82 | 0.00 | Protein coding |
| *Spata2* | ENSMUSG00000047030 | 2 | 25.88 | 0.04 | -728.17 | 0.00 | Protein coding |
| *Amh* | ENSMUSG00000035262 | 10 | 407.17 | 1.15 | -353.49 | 0.00 | Protein coding |
| *Nr4a1* | ENSMUSG00000023034 | 15 | 1336.17 | 16.00 | -83.50 | 0.00 | Protein coding |
| *Pgr* | ENSMUSG00000031870 | 9 | 332.00 | 5.42 | -61.20 | 0.00 | Protein coding |
| *Hhip* | ENSMUSG00000064325 | 8 | 13.96 | 0.23 | -60.23 | 0.00 | Protein coding |
| *Satb2* | ENSMUSG00000038331 | 1 | 47.72 | 0.80 | -59.85 | 0.00 | Protein coding |
| *Dot1l* | ENSMUSG00000061589 | 10 | 99.76 | 5.07 | -19.67 | 0.00 | Protein coding |
| *Runx2* | ENSMUSG00000039153 | 17 | 17.83 | 0.97 | -18.33 | 0.00 | Protein coding |
| *Areg* | ENSMUSG00000029378 | 5 | 4895.02 | 272.55 | -17.96 | 0.00 | Protein coding |
| *Lhcgr* | ENSMUSG00000024107 | 17 | 452.50 | 25.38 | -17.83 | 0.00 | Protein coding |
| *Elf4* | ENSMUSG00000031103 | X | 44.03 | 2.90 | -15.16 | 0.00 | Protein coding |

Chrom, chromosome; Exp, expression; TPM, transcript per million; FDR, false discovery rate; NMD, nonsense mediated decay

Table S5. Top 50 upregulated transcript isoforms in cumulus granulosa cells

| Name | ENSEMBL | Chrom | Exp in mGC (TPM) | Exp in cGC (TPM) | Fold change | FDR p-value | Biotype |
|---|---|---|---|---|---|---|---|
| *Xist-203* | ENSMUST00000320150 | X | 0.01 | 191.50 | 23203.58 | 0.00 | lncRNA |
| *Gata4-203* | ENSMUST00000121312 | 14 | 0.00 | 53.50 | 24970.38 | 0.00 | Protein coding |
| *Rack1-202* | ENSMUST00000125166 | 11 | 0.03 | 803.69 | 25082.95 | 0.00 | CDS not defined |
| *Gspt1-204* | ENSMUST00000167025 | 16 | 0.01 | 205.22 | 25297.85 | 0.00 | Protein coding |
| *Stoml2-206* | ENSMUST00000135660 | 4 | 0.01 | 173.79 | 26224.00 | 0.00 | Protein coding |
| *Bbip1-203* | ENSMUST00000235688 | 19 | 0.01 | 305.04 | 26809.20 | 0.00 | Protein coding |
| *Gnas-225* | ENSMUST00000238267 | 2 | 0.01 | 136.82 | 27000.28 | 0.00 | Protein coding |
| *Sinhcaf-209* | ENSMUST00000203164 | 6 | 0.00 | 61.55 | 27214.12 | 0.00 | Protein coding |
| *Eif4g2-207* | ENSMUST00000161079 | 7 | 0.01 | 301.93 | 27904.26 | 0.00 | CDS not defined |
| *Morf4l1-204* | ENSMUST00000187771 | 9 | 0.00 | 96.01 | 28018.85 | 0.00 | CDS not defined |
| *Akr1e1-202* | ENSMUST00000110691 | 13 | 0.00 | 46.44 | 28216.68 | 0.00 | Protein coding |
| *Cyc1-203* | ENSMUST00000229292 | 15 | 0.01 | 299.82 | 29573.82 | 0.00 | CDS not defined |
| *Rab6a-203* | ENSMUST00000107048 | 7 | 0.00 | 59.71 | 30131.88 | 0.00 | Protein coding |
| *Esd-209* | ENSMUST00000177181 | 14 | 0.01 | 195.27 | 31304.42 | 0.00 | Protein coding |
| *Tdp2-206* | ENSMUST00000225490 | 13 | 0.00 | 98.67 | 32244.52 | 0.00 | CDS not defined |
| *Cyp19a1-201* | ENSMUST00000034811 | 9 | 0.00 | 80.25 | 33754.50 | 0.00 | protein coding |
| *Eif5a-205* | ENSMUST00000108608 | 11 | 0.01 | 228.35 | 34132.27 | 0.00 | protein coding |
| *Gm49450-201* | ENSMUST00000134214 | 15 | 0.01 | 203.29 | 35973.29 | 0.00 | protein coding |
| *Snapin-205* | ENSMUST00000185005 | 3 | 0.00 | 139.91 | 36978.21 | 0.00 | NMD |
| *Gpi1-207* | ENSMUST00000205870 | 7 | 0.01 | 207.21 | 37297.13 | 0.00 | NMD |
| *Hsp90aa1-208* | ENSMUST00000155242 | 12 | 0.04 | 1633.93 | 39758.74 | 0.00 | Protein coding |
| *Rpl18-208* | ENSMUST00000211061 | 7 | 0.01 | 384.00 | 39933.35 | 0.00 | Protein coding |
| *Hnrnpk-217* | ENSMUST00000177051 | 13 | 0.01 | 249.49 | 39949.54 | 0.00 | Protein coding |
| *Csde1-216* | ENSMUST00000199571 | 3 | 0.00 | 78.88 | 40303.64 | 0.00 | Protein coding |
| *Cyb5a-205* | ENSMUST00000163083 | 18 | 0.22 | 9067.94 | 40536.81 | 0.00 | Protein coding |

| | | | | | | | |
|---|---|---|---|---|---|---|---|
| *Gnb1-208* | ENSMUST00000176637 | 4 | 0.00 | 85.94 | 42338.82 | 0.00 | Protein coding |
| *Tmem59-203* | ENSMUST00000127652 | 4 | 0.02 | 1037.42 | 46040.05 | 0.00 | CDS not defined |
| *Ahsa2-205* | ENSMUST00000128372 | 11 | 0.00 | 91.26 | 47487.07 | 0.00 | Protein coding |
| *Gabarapl2-203* | ENSMUST00000133628 | 8 | 0.01 | 346.44 | 49164.24 | 0.00 | Retained intron |
| *Bccip-204* | ENSMUST00000151711 | 7 | 0.01 | 320.00 | 49277.28 | 0.00 | CDS not defined |
| *Jpt1-202* | ENSMUST00000139980 | 11 | 0.01 | 401.50 | 49328.09 | 0.00 | CDS not defined |
| *Nudcd2-204* | ENSMUST00000141839 | 11 | 0.01 | 469.44 | 50811.85 | 0.00 | CDS not defined |
| *Tecr-214* | ENSMUST00000212990 | 8 | 0.01 | 468.90 | 53005.64 | 0.00 | Protein coding |
| *Rab1a-206* | ENSMUST00000163483 | 11 | 0.00 | 112.90 | 55664.04 | 0.00 | Protein coding |
| *Pa2g4-201* | ENSMUST00000026425 | 10 | 0.00 | 154.14 | 58569.77 | 0.00 | Protein coding |
| *Ndufa9-201* | ENSMUST00000088194 | 6 | 0.00 | 265.12 | 58915.44 | 0.00 | Protein coding |
| *Mrto4-205* | ENSMUST00000131052 | 4 | 0.01 | 387.77 | 59862.57 | 0.00 | CDS not defined |
| *Dab2-201* | ENSMUST00000078019 | 15 | 0.00 | 85.28 | 60601.20 | 0.00 | Protein coding |
| *Ranbp1-201* | ENSMUST00000052325 | 16 | 0.04 | 2799.31 | 65768.81 | 0.00 | Protein coding |
| *Tmed2-203* | ENSMUST00000135464 | 5 | 0.02 | 1411.37 | 70342.52 | 0.00 | Protein coding |
| *Rbm39-217* | ENSMUST00000146297 | 2 | 0.00 | 183.22 | 79135.28 | 0.00 | Protein coding |
| *Acly-207* | ENSMUST00000165111 | 11 | 0.00 | 110.31 | 86202.12 | 0.00 | Protein coding |
| *Vps29-201* | ENSMUST00000111729 | 5 | 0.01 | 622.74 | 90487.29 | 0.00 | NMD |
| *Ppp1ca-204* | ENSMUST00000236181 | 19 | 0.00 | 530.07 | 123727.68 | 0.00 | CDS not defined |
| *Fkbp3-205* | ENSMUST00000221166 | 12 | 0.01 | 1053.54 | 134735.85 | 0.00 | NMD |
| *Atp5po-203* | ENSMUST00000139277 | 16 | 0.01 | 1142.95 | 139636.54 | 0.00 | Protein coding |
| *Arf4-202* | ENSMUST00000112318 | 14 | 0.00 | 580.08 | 142336.35 | 0.00 | Protein coding |
| *Eif3m-202* | ENSMUST00000111110 | 2 | 0.01 | 1506.49 | 212546.89 | 0.00 | Protein coding |
| *Actg1-204* | ENSMUST00000106215 | 11 | 0.00 | 948.29 | 317548.00 | 0.00 | Protein coding |
| *Prdx2-208* | ENSMUST00000164807 | 8 | 0.00 | 1684.12 | 370134.20 | 0.00 | Protein coding |

Chrom, chromosome; Exp, expression; TPM, transcript per million; FDR, false discovery rate; NMD, nonsense mediated decay

Table S6. Top 50 downregulated transcript isoforms in cumulus granulosa cells

| Name | ENSEMBL | Chrom | Exp in mGC (TPM) | Exp in cGC (TPM) | Fold change | FDR p-value | Biotype |
|---|---|---|---|---|---|---|---|
| *Itgav-201* | ENSMUST00000028499 | 2 | 380.98 | 0.00 | -111885.58 | 0.00 | Protein coding |
| *Fat1-205* | ENSMUST00000191428 | 8 | 144.20 | 0.00 | -89746.17 | 0.00 | Protein coding |
| *Inhba-202* | ENSMUST00000164993 | 13 | 822.17 | 0.02 | -50658.23 | 0.00 | Protein coding |
| *Robo1-201* | ENSMUST00000023600 | 16 | 154.96 | 0.00 | -49070.85 | 0.00 | Protein coding |
| *Neat1-216* | ENSMUST00000301005 | 19 | 302.65 | 0.01 | -38423.55 | 0.00 | lncRNA |
| *Col4a1-201* | ENSMUST00000033898 | 8 | 143.95 | 0.00 | -36252.40 | 0.00 | Protein coding |
| *Cebpa-201* | ENSMUST00000042985 | 7 | 321.92 | 0.01 | -34914.68 | 0.00 | Protein coding |
| *Greb1-212* | ENSMUST00000162112 | 12 | 90.54 | 0.00 | -33293.86 | 0.00 | Protein coding |
| *Sulf2-201* | ENSMUST00000088086 | 2 | 197.90 | 0.01 | -32511.02 | 0.00 | Protein coding |
| *Ywhaz-204* | ENSMUST00000110362 | 15 | 362.99 | 0.01 | -32082.51 | 0.00 | Protein coding |
| *Npr1-201* | ENSMUST00000029540 | 3 | 188.77 | 0.01 | -31967.59 | 0.00 | Protein coding |
| *Peg10-201* | ENSMUST00000176204 | 6 | 111.75 | 0.00 | -30927.83 | 0.00 | Protein coding |
| *Ubr4-201* | ENSMUST00000097822 | 4 | 45.60 | 0.00 | -29347.56 | 0.00 | Protein coding |
| *Zeb2-202* | ENSMUST00000068415 | 2 | 116.20 | 0.00 | -27267.31 | 0.00 | Protein coding |
| *Snrnp70-216* | ENSMUST00000211366 | 7 | 161.04 | 0.01 | -26835.68 | 0.00 | Retained intron |
| *Ivns1abp-204* | ENSMUST00000186745 | 1 | 177.17 | 0.01 | -24416.35 | 0.00 | NMD |
| *Hnrnph1-204* | ENSMUST00000123371 | 11 | 145.20 | 0.01 | -24234.92 | 0.00 | CDS not defined |
| *Plxnb2-202* | ENSMUST00000109331 | 15 | 88.86 | 0.00 | -24110.66 | 0.00 | Protein coding |
| *Syne2-202* | ENSMUST00000085280 | 12 | 71.65 | 0.00 | -23902.14 | 0.00 | Protein coding |
| *Trrap-203* | ENSMUST00000100467 | 5 | 43.82 | 0.00 | -22856.86 | 0.00 | Protein coding |
| *Armcx2-202* | ENSMUST00000113193 | X | 146.40 | 0.01 | -22505.75 | 0.00 | Protein coding |
| *Neat1-202* | ENSMUST00000174287 | 19 | 174.15 | 0.01 | -21906.78 | 0.00 | lncRNA |
| *Psap-201* | ENSMUST00000105465 | 10 | 708.04 | 0.03 | -20793.65 | 0.00 | Protein coding |
| *Rtn4-204* | ENSMUST00000102842 | 11 | 216.01 | 0.01 | -20271.02 | 0.00 | Protein coding |
| *Nfe2l1-211* | ENSMUST00000167149 | 11 | 103.98 | 0.01 | -19990.61 | 0.00 | Protein coding |

| | | | | | | | |
|---|---|---|---|---|---|---|---|
| *Sidt2-201* | ENSMUST00000038488 | 9 | 429.27 | 0.02 | -19752.86 | 0.00 | Protein coding |
| *Fus-201* | ENSMUST00000077609 | 7 | 242.87 | 0.01 | -18584.48 | 0.00 | Protein coding |
| *AI506816-206* | ENSMUST00000244629 | 5 | 149.84 | 0.01 | -17858.84 | 0.00 | lncRNA |
| *Hnrnpk-201* | ENSMUST00000043269 | 13 | 159.70 | 0.01 | -17802.78 | 0.00 | Protein coding |
| *Itga3-201* | ENSMUST00000001548 | 11 | 75.26 | 0.00 | -17196.48 | 0.00 | Protein coding |
| *Dennd5a-201* | ENSMUST00000080437 | 7 | 78.74 | 0.00 | -16413.74 | 0.00 | Protein coding |
| *Mid1-203* | ENSMUST00000079443 | X | 646.57 | 0.04 | -15990.63 | 0.00 | Protein coding |
| *Atp13a3-202* | ENSMUST00000100013 | 16 | 45.58 | 0.00 | -15156.51 | 0.00 | Protein coding |
| *Dag1-201* | ENSMUST00000080435 | 9 | 81.73 | 0.01 | -14939.08 | 0.00 | Protein coding |
| *Csde1-211* | ENSMUST00000198180 | 3 | 92.35 | 0.01 | -14877.12 | 0.00 | Protein coding |
| *Usp34-205* | ENSMUST00000137823 | 11 | 30.89 | 0.00 | -14636.87 | 0.00 | Protein coding |
| *Acly-201* | ENSMUST00000007131 | 11 | 243.28 | 0.02 | -14239.63 | 0.00 | Protein coding |
| *Amotl2-201* | ENSMUST00000035121 | 9 | 78.01 | 0.01 | -14120.68 | 0.00 | Protein coding |
| *Mapk1ip1l-202* | ENSMUST00000166743 | 14 | 75.55 | 0.01 | -13744.25 | 0.00 | Protein coding |
| *Hyou1-206* | ENSMUST00000161318 | 9 | 82.31 | 0.01 | -13273.78 | 0.00 | Protein coding |
| *Star-202* | ENSMUST00000210565 | 8 | 493.72 | 0.04 | -12994.52 | 0.00 | Protein coding |
| *Zeb2-214* | ENSMUST00000177302 | 2 | 77.13 | 0.01 | -12881.77 | 0.00 | Protein coding |
| *Selenop-203* | ENSMUST00000159216 | 15 | 147.19 | 0.01 | -12652.54 | 0.00 | Protein coding |
| *Uba52-208* | ENSMUST00000165126 | 8 | 781.91 | 0.06 | -12611.11 | 0.00 | Protein coding |
| *Lgr4-201* | ENSMUST00000046548 | 2 | 59.32 | 0.00 | -12503.55 | 0.00 | Protein coding |
| *Midn-201* | ENSMUST00000042057 | 10 | 78.57 | 0.01 | -12195.57 | 0.00 | Protein coding |
| *Flna-201* | ENSMUST00000033699 | X | 123.45 | 0.01 | -12057.84 | 0.00 | Protein coding |
| *Mast4-207* | ENSMUST00000167058 | 13 | 27.00 | 0.00 | -11896.91 | 0.00 | Protein coding |
| *Tle3-204* | ENSMUST00000159386 | 9 | 63.07 | 0.01 | -11854.54 | 0.00 | Protein coding |
| *App-203* | ENSMUST00000226801 | 16 | 84.90 | 0.01 | -11666.87 | 0.00 | Protein coding |

Chrom, chromosome; Exp, expression; TPM, transcript per million; FDR, false discovery rate; NMD, nonsense mediated decay

Table S7: Top 50 switched-off and top 50 switched-on transcript isoforms in cGCs

| Name | ENSEMBL | Chrom | Exp in mGC (TPM) | Exp in cGC (TPM) | Fold change | FDR p-value | Biotype |
|---|---|---|---|---|---|---|---|
| *Itgav-201* | ENSMUST00000028499 | 2 | 380.98 | 0.00 | -111885.58 | 0.00 | Protein coding |
| *Fat1-205* | ENSMUST00000191428 | 8 | 144.20 | 0.00 | -89746.17 | 0.00 | Protein coding |
| *Inhba-202* | ENSMUST00000164993 | 13 | 822.17 | 0.02 | -50658.23 | 0.00 | Protein coding |
| *Robo1-201* | ENSMUST00000023600 | 16 | 154.96 | 0.00 | -49070.85 | 0.00 | Protein coding |
| *Neat1-216* | ENSMUST00000301005 | 19 | 302.65 | 0.01 | -38423.55 | 0.00 | lncRNA |
| *Col4a1-201* | ENSMUST00000033898 | 8 | 143.95 | 0.00 | -36252.40 | 0.00 | Protein coding |
| *Cebpa-201* | ENSMUST00000042985 | 7 | 321.92 | 0.01 | -34914.68 | 0.00 | Protein coding |
| *Sulf2-201* | ENSMUST00000088086 | 2 | 197.90 | 0.01 | -32511.02 | 0.00 | Protein coding |
| *Ywhaz-204* | ENSMUST00000110362 | 15 | 362.99 | 0.01 | -32082.51 | 0.00 | Protein coding |
| *Npr1-201* | ENSMUST00000029540 | 3 | 188.77 | 0.01 | -31967.59 | 0.00 | Protein coding |
| *Peg10-201* | ENSMUST00000176204 | 6 | 111.75 | 0.00 | -30927.83 | 0.00 | Protein coding |
| *Zeb2-202* | ENSMUST00000068415 | 2 | 116.20 | 0.00 | -27267.31 | 0.00 | Protein coding |
| *Snrnp70-216* | ENSMUST00000211366 | 7 | 161.04 | 0.01 | -26835.68 | 0.00 | Retained intron |
| *Ivns1abp-204* | ENSMUST00000186745 | 1 | 177.17 | 0.01 | -24416.35 | 0.00 | NMD |
| *Hnrnph1-204* | ENSMUST00000123371 | 11 | 145.20 | 0.01 | -24234.92 | 0.00 | CDS not defined |
| *Armcx2-202* | ENSMUST00000113193 | X | 146.40 | 0.01 | -22505.75 | 0.00 | Protein coding |
| *Neat1-202* | ENSMUST00000174287 | 19 | 174.15 | 0.01 | -21906.78 | 0.00 | lncRNA |
| *Psap-201* | ENSMUST00000105465 | 10 | 708.04 | 0.03 | -20793.65 | 0.00 | Protein coding |
| *Rtn4-204* | ENSMUST00000102842 | 11 | 216.01 | 0.01 | -20271.02 | 0.00 | Protein coding |
| *Nfe2l1-211* | ENSMUST00000167149 | 11 | 103.98 | 0.01 | -19990.61 | 0.00 | Protein coding |
| *Sidt2-201* | ENSMUST00000038488 | 9 | 429.27 | 0.02 | -19752.86 | 0.00 | Protein coding |
| *Fus-201* | ENSMUST00000077609 | 7 | 242.87 | 0.01 | -18584.48 | 0.00 | Protein coding |
| *AI506816-206* | ENSMUST00000244629 | 5 | 149.84 | 0.01 | -17858.84 | 0.00 | lncRNA |
| *Hnrnpk-201* | ENSMUST00000043269 | 13 | 159.70 | 0.01 | -17802.78 | 0.00 | Protein coding |
| *Mid1-203* | ENSMUST00000079443 | X | 646.57 | 0.04 | -15990.63 | 0.00 | Protein coding |
| *Acly-201* | ENSMUST00000007131 | 11 | 243.28 | 0.02 | -14239.63 | 0.00 | Protein coding |
| *Star-202* | ENSMUST00000210565 | 8 | 493.72 | 0.04 | -12994.52 | 0.00 | Protein coding |

| | | | | | | | |
|---|---|---|---|---|---|---|---|
| *Selenop-203* | ENSMUST00000159216 | 15 | 147.19 | 0.01 | -12652.54 | 0.00 | Protein coding |
| *Flna-201* | ENSMUST00000033699 | X | 123.45 | 0.01 | -12057.84 | 0.00 | Protein coding |
| *Hnrnpl-202* | ENSMUST00000172529 | 7 | 141.38 | 0.01 | -11168.57 | 0.00 | Protein coding |
| *Nap1l1-202* | ENSMUST00000217908 | 10 | 201.17 | 0.02 | -10880.62 | 0.00 | Protein coding |
| *Pigt-201* | ENSMUST00000103101 | 2 | 100.86 | 0.01 | -10810.08 | 0.00 | Protein coding |
| *Bzw1-202* | ENSMUST00000186949 | 1 | 169.19 | 0.02 | -10487.73 | 0.00 | Protein coding |
| *Rps10-208* | ENSMUST00000178774 | 17 | 492.25 | 0.05 | -10197.83 | 0.00 | Protein coding |
| *Raly-202* | ENSMUST00000058089 | 2 | 144.92 | 0.01 | -10186.52 | 0.00 | Protein coding |
| *Txnl1-201* | ENSMUST00000025476 | 18 | 111.88 | 0.01 | -9314.05 | 0.00 | Protein coding |
| *Pxdn-202* | ENSMUST00000122328 | 12 | 110.06 | 0.01 | -8665.09 | 0.00 | Protein coding |
| *Pcolce-201* | ENSMUST00000031731 | 5 | 116.90 | 0.01 | -7994.25 | 0.00 | Protein coding |
| *Tardbp-228* | ENSMUST00000190696 | 4 | 133.13 | 0.02 | -7827.56 | 0.00 | NMD |
| *Meg3-211* | ENSMUST00000166636 | 12 | 119.01 | 0.02 | -6809.19 | 0.00 | lncRNA |
| *Arpc4-204* | ENSMUST00000204802 | 6 | 121.80 | 0.03 | -4027.14 | 0.00 | Protein coding |
| *Egr3-204* | ENSMUST00000225200 | 14 | 74.97 | 0.02 | -3835.45 | 0.00 | Protein coding |
| *Srrm2-209* | ENSMUST00000190568 | 17 | 141.60 | 0.04 | -3510.28 | 0.00 | Retained intron |
| *Pim1-201* | ENSMUST00000024811 | 17 | 161.89 | 0.05 | -3418.47 | 0.00 | Protein coding |
| *Eef1g-202* | ENSMUST00000122932 | 19 | 115.39 | 0.03 | -3417.20 | 0.00 | CDS not defined |
| *Slc25a3-207* | ENSMUST00000171960 | 10 | 119.07 | 0.04 | -3378.56 | 0.00 | Retained intron |
| *Spcs2-209* | ENSMUST00000208939 | 7 | 106.68 | 0.03 | -3077.80 | 0.00 | Retained intron |
| *Gata4-201* | ENSMUST00000067417 | 14 | 43.90 | 0.04 | -1162.28 | 0.00 | Protein coding |
| *Prmt1-205* | ENSMUST00000207659 | 7 | 21.31 | 0.03 | -839.03 | 0.00 | Protein coding |
| *Foxp4-204* | ENSMUST00000113265 | 17 | 16.68 | 0.02 | -783.60 | 0.00 | Protein coding |
| *Prelid3b-202* | ENSMUST00000117442 | 2 | 0.05 | 200.11 | 4191.98 | 0.00 | Protein coding |
| *Timm13-204* | ENSMUST00000219896 | 10 | 0.04 | 184.04 | 5034.20 | 0.00 | Protein coding |
| *Hax1-204* | ENSMUST00000197725 | 3 | 0.04 | 214.18 | 5037.84 | 0.00 | Protein coding |
| *Psma6-202* | ENSMUST00000162711 | 12 | 0.04 | 232.29 | 5193.18 | 0.00 | NMD |
| *Psme2-206* | ENSMUST00000161027 | 14 | 0.05 | 329.80 | 6780.33 | 0.00 | Retained intron |
| *Malat1-205* | ENSMUST00000245150 | 19 | 0.03 | 219.62 | 6838.74 | 0.00 | lncRNA |
| *Hnrnpf-210* | ENSMUST00000180341 | 6 | 0.04 | 331.69 | 7982.86 | 0.00 | Protein coding |
| *Psmc3-203* | ENSMUST00000111441 | 2 | 0.02 | 171.75 | 8168.37 | 0.00 | Protein coding |

| | | | | | | | |
|---|---|---|---|---|---|---|---|
| *Rrm1-202* | ENSMUST00000209630 | 7 | 0.01 | 70.92 | 9968.81 | 0.00 | NMD |
| *Crem-220* | ENSMUST00000140912 | 18 | 0.02 | 294.71 | 14068.19 | 0.00 | CDS not defined |
| *Creg1-201* | ENSMUST00000040298 | 1 | 0.01 | 221.78 | 18112.93 | 0.00 | Protein coding |
| *Ssb-202* | ENSMUST00000112260 | 2 | 0.01 | 176.16 | 20322.71 | 0.00 | Protein coding |
| *Sumo3-203* | ENSMUST00000124024 | 10 | 0.01 | 166.95 | 22458.31 | 0.00 | Protein coding |
| *Xist-203* | ENSMUST00000320150 | X | 0.01 | 191.50 | 23203.58 | 0.00 | lncRNA |
| *Smc2-203* | ENSMUST00000131163 | 4 | 0.01 | 192.55 | 24044.48 | 0.00 | CDS not defined |
| *Gata4-203* | ENSMUST00000121312 | 14 | 0.00 | 53.50 | 24970.38 | 0.00 | Protein coding |
| *Rack1-202* | ENSMUST00000125166 | 11 | 0.03 | 803.69 | 25082.95 | 0.00 | CDS not defined |
| *Gspt1-204* | ENSMUST00000167025 | 16 | 0.01 | 205.22 | 25297.85 | 0.00 | Protein coding |
| *Stoml2-206* | ENSMUST00000135660 | 4 | 0.01 | 173.79 | 26224.00 | 0.00 | Protein coding |
| *Bbip1-203* | ENSMUST00000235688 | 19 | 0.01 | 305.04 | 26809.20 | 0.00 | Protein coding |
| *Eif4g2-207* | ENSMUST00000161079 | 7 | 0.01 | 301.93 | 27904.26 | 0.00 | CDS not defined |
| *Cyc1-203* | ENSMUST00000229292 | 15 | 0.01 | 299.82 | 29573.82 | 0.00 | CDS not defined |
| *Esd-209* | ENSMUST00000177181 | 14 | 0.01 | 195.27 | 31304.42 | 0.00 | Protein coding |
| *Cyp19a1-201* | ENSMUST00000034811 | 9 | 0.00 | 80.25 | 33754.50 | 0.00 | Protein coding |
| *Eif5a-205* | ENSMUST00000108608 | 11 | 0.01 | 228.35 | 34132.27 | 0.00 | Protein coding |
| *Gm49450-201* | ENSMUST00000134214 | 15 | 0.01 | 203.29 | 35973.29 | 0.00 | Protein coding |
| *Gpi1-207* | ENSMUST00000205870 | 7 | 0.01 | 207.21 | 37297.13 | 0.00 | NMD |
| *Hsp90aa1-208* | ENSMUST00000155242 | 12 | 0.04 | 1633.93 | 39758.74 | 0.00 | Protein coding |
| *Rpl18-208* | ENSMUST00000211061 | 7 | 0.01 | 384.00 | 39933.35 | 0.00 | Protein coding |
| *Hnrnpk-217* | ENSMUST00000177051 | 13 | 0.01 | 249.49 | 39949.54 | 0.00 | Retained intron |
| *Tmem59-203* | ENSMUST00000127652 | 4 | 0.02 | 1037.42 | 46040.05 | 0.00 | CDS not defined |
| *Gabarapl2-203* | ENSMUST00000133628 | 8 | 0.01 | 346.44 | 49164.24 | 0.00 | Retained intron |
| *Bccip-204* | ENSMUST00000151711 | 7 | 0.01 | 320.00 | 49277.28 | 0.00 | CDS not defined |
| *Jpt1-202* | ENSMUST00000139980 | 11 | 0.01 | 401.50 | 49328.09 | 0.00 | CDS not defined |
| *Idh3g-202* | ENSMUST00000129070 | X | 0.01 | 319.27 | 49959.55 | 0.00 | Retained intron |
| *Nudcd2-204* | ENSMUST00000141839 | 11 | 0.01 | 469.44 | 50811.85 | 0.00 | CDS not defined |
| *Tecr-214* | ENSMUST00000212990 | 8 | 0.01 | 468.90 | 53005.64 | 0.00 | Protein coding |
| *Ndufa9-201* | ENSMUST00000088194 | 6 | 0.00 | 265.12 | 58915.44 | 0.00 | Protein coding |
| *Mrto4-205* | ENSMUST00000131052 | 4 | 0.01 | 387.77 | 59862.57 | 0.00 | CDS not defined |

| | | | | | | | |
|---|---|---|---|---|---|---|---|
| *Ranbp1-201* | ENSMUST00000052325 | 16 | 0.04 | 2799.31 | 65768.81 | 0.00 | Protein coding |
| *Tmed2-203* | ENSMUST00000135464 | 5 | 0.02 | 1411.37 | 70342.52 | 0.00 | Protein coding |
| *Rbm39-217* | ENSMUST00000146297 | 2 | 0.00 | 183.22 | 79135.28 | 0.00 | Protein coding |
| *Vps29-201* | ENSMUST00000111729 | 5 | 0.01 | 622.74 | 90487.29 | 0.00 | NMD |
| *Ppp1ca-204* | ENSMUST00000236181 | 19 | 0.00 | 530.07 | 123727.68 | 0.00 | CDS not defined |
| *Fkbp3-205* | ENSMUST00000221166 | 12 | 0.01 | 1053.54 | 134735.85 | 0.00 | NMD |
| *Atp5po-203* | ENSMUST00000139277 | 16 | 0.01 | 1142.95 | 139636.54 | 0.00 | Protein coding |
| *Arf4-202* | ENSMUST00000112318 | 14 | 0.00 | 580.08 | 142336.35 | 0.00 | Protein coding |
| *Eif3m-202* | ENSMUST00000111110 | 2 | 0.01 | 1506.49 | 212546.89 | 0.00 | Protein coding |
| *Actg1-204* | ENSMUST00000106215 | 11 | 0.00 | 948.29 | 317548.00 | 0.00 | Protein coding |
| *Prdx2-208* | ENSMUST00000164807 | 8 | 0.00 | 1684.12 | 370134.20 | 0.00 | Protein coding |

Chrom, chromosome; Exp, expression; TPM, transcript per million; FDR, false discovery rate; NMD, nonsense mediated decay

Table S8. Differentially expressed transcript isoforms of genes, with one isoform switched-off and another switched-on

| Name | ENSEMBL | Chrom | Exp in mGC (TPM) | Exp in cGC (TPM) | Fold change | FDR p-value | Biotype |
|---|---|---|---|---|---|---|---|
| *Actg1-204* | ENSMUST00000106215 | 11 | 0.00 | 948.29 | 317548.00 | 0.00 | Protein coding |
| *Rbm39-217* | ENSMUST00000146297 | 2 | 0.00 | 183.22 | 79135.28 | 0.00 | Protein coding |
| *Gnb1-208* | ENSMUST00000176637 | 4 | 0.00 | 85.94 | 42338.82 | 0.00 | Protein coding |
| *Csde1-216* | ENSMUST00000199571 | 3 | 0.00 | 78.88 | 40303.64 | 0.00 | Protein coding |
| *Rab6a-203* | ENSMUST00000107048 | 7 | 0.00 | 59.71 | 30131.88 | 0.00 | Protein coding |
| *Gnas-225* | ENSMUST00000238267 | 2 | 0.01 | 136.82 | 27000.28 | 0.00 | Protein coding |
| *Gata4-203* | ENSMUST00000121312 | 14 | 0.00 | 53.50 | 24970.38 | 0.00 | Protein coding |
| *Macf1-211* | ENSMUST00000149022 | 4 | 0.00 | 57.53 | 24126.50 | 0.00 | Protein coding |
| *Cdv3-207* | ENSMUST00000190226 | 9 | 0.00 | 38.77 | 22816.85 | 0.00 | Protein coding |
| *Gps1-205* | ENSMUST00000153678 | 11 | 0.00 | 72.40 | 20457.36 | 0.00 | NMD |
| *Lman1-205* | ENSMUST00000150133 | 18 | 0.01 | 134.25 | 18924.00 | 0.00 | CDS not defined |
| *Neu3-203* | ENSMUST00000250003 | 7 | 0.00 | 30.24 | 17612.04 | 0.00 | Protein coding |
| *Crem-220* | ENSMUST00000140912 | 18 | 0.02 | 294.71 | 14068.19 | 0.00 | CDS not defined |
| *Nars1-209* | ENSMUST00000236873 | 18 | 0.00 | 29.51 | 13472.19 | 0.00 | NMD |
| *Pja2-202* | ENSMUST00000024889 | 17 | 0.00 | 17.32 | 12690.45 | 0.00 | Protein coding |
| *Gtf2i-207* | ENSMUST00000172904 | 5 | 0.00 | 20.77 | 12033.78 | 0.00 | CDS not defined |
| *Atad1-202* | ENSMUST00000235142 | 19 | 0.00 | 28.38 | 11787.80 | 0.00 | Protein coding |
| *Setdb1-203* | ENSMUST00000107171 | 3 | 0.00 | 12.60 | 10766.89 | 0.00 | Protein coding |
| *Flvcr1-205* | ENSMUST00000194589 | 1 | 0.00 | 22.46 | 10615.58 | 0.00 | retained_intron |
| *Ndfip2-203* | ENSMUST00000149358 | 14 | 0.00 | 35.03 | 9565.63 | 0.00 | CDS not defined |
| *Gramd1b-207* | ENSMUST00000137454 | 9 | 0.00 | 14.77 | 7155.39 | 0.00 | Protein coding |
| *Fbxo11-206* | ENSMUST00000235112 | 17 | 0.00 | 13.57 | 6345.30 | 0.00 | Protein coding |
| *Sgce-207* | ENSMUST00000126151 | 6 | 0.00 | 22.60 | 6261.06 | 0.00 | Protein coding |
| *Rsrc2-205* | ENSMUST00000182015 | 5 | 0.01 | 33.43 | 5493.72 | 0.00 | NMD |
| *Ppp1cc-206* | ENSMUST00000134719 | 5 | 0.01 | 31.13 | 4699.18 | 0.00 | NMD |
| *Cmip-201* | ENSMUST00000095172 | 8 | 0.00 | 9.35 | 3633.05 | 0.00 | Protein coding |
| *Cndp2-207* | ENSMUST00000236415 | 18 | 0.01 | 48.84 | 3345.78 | 0.00 | CDS not defined |

| | | | | | | | |
|---|---|---|---|---|---|---|---|
| *Mprip-209* | ENSMUST00000156111 | 11 | 0.01 | 37.79 | 2629.07 | 0.00 | Protein coding |
| *Map4k5-201* | ENSMUST00000110567 | 12 | 0.00 | 12.33 | 2599.56 | 0.00 | Protein coding |
| *Hspa8-209* | ENSMUST00000153847 | 9 | 0.30 | 568.27 | 1881.04 | 0.00 | CDS not defined |
| *Tor1b-206* | ENSMUST00000156711 | 2 | 0.01 | 16.89 | 1849.60 | 0.00 | NMD |
| *Add1-213* | ENSMUST00000152805 | 5 | 0.01 | 21.01 | 1491.15 | 0.00 | Protein coding |
| *Glce-201* | ENSMUST00000034785 | 9 | 0.01 | 10.37 | 1441.33 | 0.00 | Protein coding |
| *Arpc1b-205* | ENSMUST00000136074 | 5 | 0.01 | 12.06 | 1389.12 | 0.00 | Protein coding |
| *Gnb2-209* | ENSMUST00000143495 | 5 | 0.00 | 6.51 | 1364.69 | 0.00 | Protein coding |
| *Stk3-205* | ENSMUST00000226555 | 15 | 0.01 | 12.72 | 1160.84 | 0.00 | Protein coding |
| *Ezh2-203* | ENSMUST00000114616 | 6 | 0.01 | 13.52 | 1117.91 | 0.00 | Protein coding |
| *Alg1-201* | ENSMUST00000049207 | 16 | 0.02 | 16.59 | 799.25 | 0.00 | Protein coding |
| *Vps35l-214* | ENSMUST00000176197 | 7 | 0.05 | 20.68 | 418.66 | 0.00 | retained_intron |
| *Anp32e-208* | ENSMUST00000170213 | 3 | 0.07 | 29.53 | 394.93 | 0.00 | NMD |
| *Zeb2-213* | ENSMUST00000176732 | 2 | 0.09 | 20.82 | 224.19 | 0.00 | Protein coding |
| *Neat1-219* | ENSMUST00000301008 | 19 | 0.09 | 12.10 | 141.19 | 0.00 | lncRNA |
| *Esr2-203* | ENSMUST00000110421 | 12 | 0.25 | 20.67 | 81.50 | 0.00 | Protein coding |
| *Ss18-210* | ENSMUST00000235011 | 18 | 0.31 | 23.45 | 75.88 | 0.00 | Protein coding |
| *Tnpo1-201* | ENSMUST00000109399 | 13 | 0.22 | 15.98 | 72.46 | 0.00 | Protein coding |
| *Rbm6-202* | ENSMUST00000181986 | 9 | 0.34 | 17.43 | 51.08 | 0.00 | NMD |
| *Ilf3-202* | ENSMUST00000115414 | 9 | 1.35 | 11.49 | 8.48 | 0.00 | Protein coding |
| *Elovl1-203* | ENSMUST00000102673 | 4 | 3.66 | 14.12 | 3.86 | 0.00 | Protein coding |
| *Stk3-201* | ENSMUST00000018476 | 15 | 16.55 | 0.73 | -22.57 | 0.00 | Protein coding |
| *Crem-239* | ENSMUST00000165086 | 18 | 18.11 | 0.77 | -23.57 | 0.00 | Protein coding |
| *Cdv3-203* | ENSMUST00000116517 | 9 | 18.73 | 0.17 | -107.87 | 0.00 | Protein coding |
| *Hspa8-207* | ENSMUST00000140984 | 9 | 18.25 | 0.13 | -137.33 | 0.00 | retained_intron |
| *Vps35l-204* | ENSMUST00000106553 | 7 | 15.99 | 0.04 | -410.07 | 0.00 | Protein coding |
| *Tor1b-203* | ENSMUST00000133544 | 2 | 11.65 | 0.03 | -417.73 | 0.00 | Protein coding |
| *Rbm39-201* | ENSMUST00000029149 | 2 | 145.44 | 0.33 | -440.16 | 0.00 | Protein coding |
| *Setdb1-204* | ENSMUST00000124638 | 3 | 10.40 | 0.02 | -462.94 | 0.00 | retained_intron |
| *Ppp1cc-201* | ENSMUST00000086294 | 5 | 33.95 | 0.07 | -501.42 | 0.00 | Protein coding |

| | | | | | | | |
|---|---|---|---|---|---|---|---|
| *Arpc1b-202* | ENSMUST00000126343 | 5 | 14.40 | 0.03 | -549.68 | 0.00 | retained_intron |
| *Alg1-202* | ENSMUST00000100196 | 16 | 32.49 | 0.05 | -664.00 | 0.00 | Protein coding |
| *Rsrc2-202* | ENSMUST00000057795 | 5 | 24.08 | 0.04 | -679.17 | 0.00 | Protein coding |
| *Flvcr1-201* | ENSMUST00000085635 | 1 | 29.14 | 0.03 | -921.41 | 0.00 | Protein coding |
| *Ilf3-201* | ENSMUST00000067646 | 9 | 35.01 | 0.04 | -932.32 | 0.00 | Protein coding |
| *Anp32e-203* | ENSMUST00000167876 | 3 | 32.35 | 0.03 | -955.47 | 0.00 | Protein coding |
| *Fbxo11-201* | ENSMUST00000005504 | 17 | 24.80 | 0.03 | -959.23 | 0.00 | Protein coding |
| *Rbm6-207* | ENSMUST00000183032 | 9 | 19.73 | 0.02 | -962.00 | 0.00 | Protein coding |
| *Gnb2-212* | ENSMUST00000150063 | 5 | 16.50 | 0.02 | -1071.82 | 0.00 | Protein coding |
| *Actg1-205* | ENSMUST00000128055 | 11 | 27.38 | 0.02 | -1151.94 | 0.00 | Protein coding |
| *Gata4-201* | ENSMUST00000067417 | 14 | 43.90 | 0.04 | -1162.28 | 0.00 | Protein coding |
| *Lman1-202* | ENSMUST00000120461 | 18 | 21.06 | 0.02 | -1245.56 | 0.00 | Protein coding |
| *Nars1-205* | ENSMUST00000236186 | 18 | 17.31 | 0.01 | -1288.18 | 0.00 | Protein coding |
| *Tnpo1-204* | ENSMUST00000179301 | 13 | 13.18 | 0.01 | -1388.95 | 0.00 | Protein coding |
| *Ezh2-201* | ENSMUST00000081721 | 6 | 12.26 | 0.01 | -1432.87 | 0.00 | Protein coding |
| *Ndfip2-206* | ENSMUST00000181969 | 14 | 16.85 | 0.01 | -1622.06 | 0.00 | Protein coding |
| *Rab6a-202* | ENSMUST00000098252 | 7 | 14.85 | 0.01 | -1923.59 | 0.00 | Protein coding |
| *Add1-204* | ENSMUST00000114338 | 5 | 12.24 | 0.01 | -1958.81 | 0.00 | Protein coding |
| *Ss18-201* | ENSMUST00000040924 | 18 | 17.70 | 0.01 | -2284.46 | 0.00 | Protein coding |
| *Cndp2-202* | ENSMUST00000168419 | 18 | 28.05 | 0.01 | -2413.02 | 0.00 | Protein coding |
| *Pja2-203* | ENSMUST00000172733 | 17 | 13.01 | 0.00 | -2648.22 | 0.00 | Protein coding |
| *Elovl1-207* | ENSMUST00000167636 | 4 | 55.42 | 0.02 | -3101.47 | 0.00 | Protein coding |
| *Sgce-205* | ENSMUST00000115579 | 6 | 55.30 | 0.01 | -3908.28 | 0.00 | Protein coding |
| *Map4k5-202* | ENSMUST00000110570 | 12 | 21.31 | 0.01 | -3991.65 | 0.00 | Protein coding |
| *Neu3-201* | ENSMUST00000036331 | 7 | 29.41 | 0.01 | -4066.88 | 0.00 | Protein coding |
| *Gps1-202* | ENSMUST00000116305 | 11 | 55.96 | 0.01 | -4264.29 | 0.00 | Protein coding |
| *Gnas-202* | ENSMUST00000087871 | 2 | 29.97 | 0.01 | -4807.40 | 0.00 | Protein coding |
| *Cmip-202* | ENSMUST00000166750 | 8 | 58.44 | 0.01 | -5859.16 | 0.00 | Protein coding |
| *Esr2-201* | ENSMUST00000076634 | 12 | 43.37 | 0.01 | -5916.31 | 0.00 | Protein coding |
| *Gtf2i-203* | ENSMUST00000111261 | 5 | 27.12 | 0.00 | -5930.85 | 0.00 | Protein coding |

| | | | | | | | |
|---|---|---|---|---|---|---|---|
| *Atad1-201* | ENSMUST00000070210 | 19 | 55.34 | 0.01 | -6171.28 | 0.00 | Protein coding |
| *Gnb1-203* | ENSMUST00000165335 | 4 | 48.57 | 0.01 | -6358.84 | 0.00 | Protein coding |
| *Glce-203* | ENSMUST00000185675 | 9 | 34.38 | 0.00 | -6990.33 | 0.00 | Protein coding |
| *Zeb2-201* | ENSMUST00000028229 | 2 | 36.17 | 0.00 | -8040.69 | 0.00 | Protein coding |
| *Mprip-203* | ENSMUST00000108751 | 11 | 54.47 | 0.01 | -9098.96 | 0.00 | Protein coding |
| *Gramd1b-212* | ENSMUST00000165104 | 9 | 64.36 | 0.01 | -10328.83 | 0.00 | Protein coding |
| *Macf1-206* | ENSMUST00000134458 | 4 | 18.28 | 0.00 | -10662.16 | 0.00 | Protein coding |
| *Csde1-211* | ENSMUST00000198180 | 3 | 92.35 | 0.01 | -14877.12 | 0.00 | Protein coding |
| *Neat1-216* | ENSMUST00000301005 | 19 | 302.65 | 0.01 | -38423.55 | 0.00 | lncRNA |

Chrom, chromosome; Exp, expression; TPM, transcript per million; FDR, false discovery rate; NMD, nonsense mediated decay

Table S9: Top 50 differentially expressed transcript isoforms of epigenetic regulators

| *Name* | ENSEMBL | Chrom | Exp in mGC (TPM) | Exp in cGC (TPM) | Fold change | FDR p-value | Biotype |
|---|---|---|---|---|---|---|---|
| *Ywhaz-204* | ENSMUST00000110362 | 15 | 362.99 | 0.01 | -32082.51 | 0.00 | Protein coding |
| *Trrap-203* | ENSMUST00000100467 | 5 | 43.82 | 0.00 | -22856.86 | 0.00 | Protein coding |
| *Nap1l1-202* | ENSMUST00000217908 | 10 | 201.17 | 0.02 | -10880.62 | 0.00 | Protein coding |
| *Aebp2-202* | ENSMUST00000087614 | 6 | 47.85 | 0.01 | -7382.34 | 0.00 | Protein coding |
| *Men1-201* | ENSMUST00000056391 | 19 | 47.00 | 0.01 | -5853.22 | 0.00 | Protein coding |
| *Brd2-202* | ENSMUST00000095347 | 17 | 98.48 | 0.03 | -3601.02 | 0.00 | Protein coding |
| *Brd2-207* | ENSMUST00000154232 | 17 | 64.87 | 0.02 | -3457.68 | 0.00 | NMD |
| *Ogt-203* | ENSMUST00000147635 | X | 57.43 | 0.04 | -1582.63 | 0.00 | CDS not defined |
| *Prmt1-202* | ENSMUST00000107843 | 7 | 39.18 | 0.06 | -615.69 | 0.00 | Protein coding |
| *Morf4l1-202* | ENSMUST00000169860 | 9 | 320.35 | 0.59 | -540.10 | 0.00 | Protein coding |
| *Atn1-201* | ENSMUST00000088357 | 6 | 46.98 | 0.16 | -300.66 | 0.00 | Protein coding |
| *Cdc73-206* | ENSMUST00000212510 | 1 | 57.54 | 0.40 | -144.92 | 0.00 | CDS not defined |
| *Dot1l-201* | ENSMUST00000105336 | 10 | 75.91 | 1.06 | -71.31 | 0.00 | Protein coding |
| *Npm1-203* | ENSMUST00000101375 | 11 | 169.85 | 2.75 | -61.72 | 0.00 | Protein coding |
| *Ctbp1-211* | ENSMUST00000202868 | 5 | 64.15 | 1.10 | -58.46 | 0.00 | Protein coding |
| *Ogt-204* | ENSMUST00000150161 | X | 75.12 | 1.90 | -39.63 | 0.00 | Retained intron |
| *Usp3-205* | ENSMUST00000127569 | 9 | 108.16 | 2.79 | -38.73 | 0.00 | Protein coding |
| *Fbl-201* | ENSMUST00000042405 | 7 | 167.32 | 4.97 | -33.69 | 0.00 | Protein coding |
| *Mta1-202* | ENSMUST00000069690 | 12 | 48.83 | 1.59 | -30.64 | 0.00 | Protein coding |
| *Hmgb1-203* | ENSMUST00000110505 | 5 | 197.61 | 8.29 | -23.83 | 0.00 | Protein coding |
| *Bap1-201* | ENSMUST00000022458 | 14 | 46.24 | 2.16 | -21.42 | 0.00 | Protein coding |
| *Sfpq-204* | ENSMUST00000143168 | 4 | 59.95 | 3.22 | -18.59 | 0.00 | CDS not defined |
| *Kdm6b-201* | ENSMUST00000094077 | 11 | 53.02 | 3.11 | -17.04 | 0.00 | Protein coding |
| *Chd4-203* | ENSMUST00000112392 | 6 | 57.53 | 3.53 | -16.31 | 0.00 | Protein coding |
| *Wsb2-201* | ENSMUST00000031309 | 5 | 108.48 | 7.36 | -14.74 | 0.00 | Protein coding |
| *Morf4l2-204* | ENSMUST00000113097 | X | 5.69 | 17.47 | 3.07 | 0.01 | Protein coding |
| *Hmgn2-203* | ENSMUST00000102553 | 4 | 193.54 | 1823.14 | 9.42 | 0.00 | Protein coding |

| | | | | | | | |
|---|---|---|---|---|---|---|---|
| *Ube2d3-201* | ENSMUST00000106291 | 3 | 58.85 | 610.83 | 10.38 | 0.00 | Protein coding |
| *Srsf3-202* | ENSMUST00000130216 | 17 | 31.37 | 334.13 | 10.65 | 0.00 | Protein coding |
| *Gadd45g-201* | ENSMUST00000021903 | 13 | 20.50 | 221.41 | 10.80 | 0.00 | Protein coding |
| *Sin3b-202* | ENSMUST00000109950 | 8 | 42.54 | 472.20 | 11.10 | 0.00 | Protein coding |
| *Dpy30-206* | ENSMUST00000233581 | 17 | 15.75 | 197.19 | 12.52 | 0.00 | Protein coding |
| *Dnajc2-201* | ENSMUST00000030771 | 5 | 11.92 | 170.42 | 14.30 | 0.00 | Protein coding |
| *Skp1-201* | ENSMUST00000037324 | 11 | 95.87 | 1412.21 | 14.73 | 0.00 | Protein coding |
| *Aurka-203* | ENSMUST00000109140 | 2 | 8.51 | 134.39 | 15.80 | 0.00 | Protein coding |
| *Exosc3-201* | ENSMUST00000030003 | 4 | 14.91 | 238.66 | 16.01 | 0.00 | Protein coding |
| *Exosc8-201* | ENSMUST00000029316 | 3 | 8.97 | 158.13 | 17.62 | 0.00 | Protein coding |
| *Clns1a-201* | ENSMUST00000026506 | 7 | 21.76 | 439.95 | 20.22 | 0.00 | Protein coding |
| *Pbk-201* | ENSMUST00000022612 | 14 | 12.36 | 267.37 | 21.63 | 0.00 | Protein coding |
| *Hat1-201* | ENSMUST00000028408 | 2 | 25.90 | 574.73 | 22.19 | 0.00 | Protein coding |
| *Exosc9-201* | ENSMUST00000029269 | 3 | 7.90 | 185.03 | 23.42 | 0.00 | Protein coding |
| *Actr6-201* | ENSMUST00000020109 | 10 | 3.01 | 132.79 | 44.16 | 0.00 | Protein coding |
| *Exosc7-201* | ENSMUST00000026891 | 9 | 6.77 | 343.42 | 50.75 | 0.00 | Protein coding |
| *Hmgb1-202* | ENSMUST00000093196 | 5 | 11.78 | 711.56 | 60.38 | 0.00 | Protein coding |
| *Rbbp7-202* | ENSMUST00000112326 | X | 16.76 | 1159.36 | 69.16 | 0.00 | Protein coding |
| *Asf1a-204* | ENSMUST00000219271 | 10 | 4.76 | 594.12 | 124.76 | 0.00 | Protein coding |
| *Ube2d3-207* | ENSMUST00000196878 | 3 | 2.32 | 342.90 | 147.64 | 0.00 | Retained intron |
| *Dpy30-203* | ENSMUST00000232687 | 17 | 1.69 | 371.23 | 219.08 | 0.00 | Protein coding |
| *Morf4l2-213* | ENSMUST00000166478 | X | 1.51 | 699.38 | 463.50 | 0.00 | Protein coding |
| *Morf4l1-204* | ENSMUST00000187771 | 9 | 0.00 | 96.01 | 28018.85 | 0.00 | CDS not defined |

Chrom, chromosome; Exp, expression; TPM, transcript per million; FDR, false discovery rate; NMD, nonsense mediated decay

Table S10: Top 50 differentially expressed transcript isoforms of transcription factors

| Name | ENSEMBL | Chrom | Exp in mGC (TPM) | Exp in cGC (TPM) | Fold change | FDR p-value | Biotype |
|---|---|---|---|---|---|---|---|
| *Cebpa-201* | ENSMUST00000042985 | 7 | 321.92 | 0.01 | -34914.68 | 0.00 | Protein coding |
| *Fus-201* | ENSMUST00000077609 | 7 | 242.87 | 0.01 | -18584.48 | 0.00 | Protein coding |
| *Mid1-203* | ENSMUST00000079443 | X | 646.57 | 0.04 | -15990.63 | 0.00 | Protein coding |
| *Pim1-201* | ENSMUST00000024811 | 17 | 161.89 | 0.05 | -3418.47 | 0.00 | Protein coding |
| *Pgr-203* | ENSMUST00000151080 | 9 | 200.93 | 0.21 | -962.23 | 0.00 | retained intron |
| *Morf4l1-202* | ENSMUST00000169860 | 9 | 320.35 | 0.59 | -540.10 | 0.00 | Protein coding |
| *Jund-201* | ENSMUST00000095267 | 8 | 354.28 | 1.54 | -229.60 | 0.00 | Protein coding |
| *Ctnnb1-201* | ENSMUST00000007130 | 9 | 385.25 | 2.43 | -158.35 | 0.00 | Protein coding |
| *Zyx-207* | ENSMUST00000203652 | 6 | 164.92 | 1.08 | -152.43 | 0.00 | Protein coding |
| *Nr5a2-202* | ENSMUST00000168126 | 1 | 351.52 | 3.52 | -99.86 | 0.00 | Protein coding |
| *Nr4a1-201* | ENSMUST00000023779 | 15 | 1196.63 | 14.59 | -81.99 | 0.00 | Protein coding |
| *Sfrp4-203* | ENSMUST00000222464 | 13 | 366.20 | 4.71 | -77.78 | 0.00 | Protein coding |
| *Jun-201* | ENSMUST00000107094 | 4 | 167.85 | 3.97 | -42.24 | 0.00 | Protein coding |
| *S100a9-201* | ENSMUST00000069960 | 3 | 217.89 | 5.19 | -41.99 | 0.00 | Protein coding |
| *Jun-202* | ENSMUST00000249436 | 4 | 169.62 | 4.69 | -36.15 | 0.00 | Protein coding |
| *Junb-201* | ENSMUST00000064922 | 8 | 889.68 | 27.80 | -32.00 | 0.00 | Protein coding |
| *Sfrp4-201* | ENSMUST00000002883 | 13 | 1144.59 | 47.97 | -23.86 | 0.00 | Protein coding |
| *Hmgb1-203* | ENSMUST00000110505 | 5 | 197.61 | 8.29 | -23.83 | 0.00 | Protein coding |
| *Enc1-201* | ENSMUST00000041623 | 13 | 838.54 | 45.85 | -18.29 | 0.00 | Protein coding |
| *Hnrnpa2b1-208* | ENSMUST00000204158 | 6 | 330.38 | 20.15 | -16.40 | 0.00 | NMD |
| *Btg1-201* | ENSMUST00000038377 | 10 | 625.84 | 39.74 | -15.75 | 0.00 | Protein coding |
| *Gnas-208* | ENSMUST00000109087 | 2 | 768.53 | 49.45 | -15.54 | 0.00 | Protein coding |
| *Cited2-201* | ENSMUST00000038107 | 10 | 945.97 | 67.47 | -14.02 | 0.00 | Protein coding |
| *Eif3k-201* | ENSMUST00000066070 | 7 | 212.03 | 17.91 | -11.84 | 0.00 | Protein coding |
| *Ddx5-208* | ENSMUST00000133426 | 11 | 419.64 | 42.95 | -9.77 | 0.00 | NMD |
| *Btf3-205* | ENSMUST00000152704 | 13 | 188.45 | 1189.14 | 6.31 | 0.00 | Protein coding |
| *Sumo2-207* | ENSMUST00000153892 | 11 | 93.47 | 664.57 | 7.11 | 0.00 | Protein coding |

| | | | | | | | |
|---|---|---|---|---|---|---|---|
| *Hmgn2-203* | ENSMUST00000102553 | 4 | 193.54 | 1823.14 | 9.42 | 0.00 | Protein coding |
| *Ubb-201* | ENSMUST00000019649 | 11 | 580.27 | 5506.73 | 9.49 | 0.00 | Protein coding |
| *Myef2-206* | ENSMUST00000142718 | 2 | 12.89 | 139.87 | 10.85 | 0.00 | Protein coding |
| *Id3-201* | ENSMUST00000008016 | 4 | 189.91 | 2370.09 | 12.48 | 0.00 | Protein coding |
| *Naca-201* | ENSMUST00000073868 | 10 | 251.19 | 3323.30 | 13.23 | 0.00 | Protein coding |
| *Park7-201* | ENSMUST00000030805 | 4 | 49.25 | 682.05 | 13.85 | 0.00 | Protein coding |
| *Sub1-202* | ENSMUST00000110504 | 15 | 65.15 | 939.43 | 14.42 | 0.00 | Protein coding |
| *H2az1-201* | ENSMUST00000041045 | 3 | 574.11 | 8795.43 | 15.32 | 0.00 | Protein coding |
| *Tceal8-201* | ENSMUST00000060101 | X | 45.65 | 845.36 | 18.52 | 0.00 | Protein coding |
| *Cnbp-203* | ENSMUST00000113619 | 6 | 83.34 | 1757.71 | 21.09 | 0.00 | Protein coding |
| *Oaz1-206* | ENSMUST00000180036 | 10 | 147.94 | 3299.02 | 22.30 | 0.00 | Protein coding |
| *Atp5f1c-203* | ENSMUST00000114897 | 2 | 74.92 | 1698.51 | 22.67 | 0.00 | Protein coding |
| *Rgcc-201* | ENSMUST00000022595 | 14 | 178.06 | 4038.47 | 22.68 | 0.00 | Protein coding |
| *Hmgb2-201* | ENSMUST00000067925 | 8 | 47.94 | 1134.21 | 23.66 | 0.00 | Protein coding |
| *Bud31-207* | ENSMUST00000162594 | 5 | 33.40 | 879.74 | 26.34 | 0.00 | Protein coding |
| *Eloc-201* | ENSMUST00000115352 | 1 | 27.30 | 874.13 | 32.02 | 0.00 | Protein coding |
| *Rps27a-202* | ENSMUST00000102845 | 11 | 21.80 | 930.06 | 42.67 | 0.00 | Protein coding |
| *Sumo1-201* | ENSMUST00000091374 | 1 | 42.60 | 2211.55 | 51.91 | 0.00 | Protein coding |
| *Gtf2a2-202* | ENSMUST00000118198 | 9 | 15.64 | 1002.79 | 64.12 | 0.00 | Protein coding |
| *Rbbp7-202* | ENSMUST00000112326 | X | 16.76 | 1159.36 | 69.16 | 0.00 | Protein coding |
| *Crabp2-201* | ENSMUST00000005019 | 3 | 15.84 | 1137.91 | 71.85 | 0.00 | Protein coding |
| *Rpf2-206* | ENSMUST00000183309 | 10 | 9.86 | 754.24 | 76.48 | 0.00 | Protein coding |
| *Hspa8-202* | ENSMUST00000117557 | 9 | 12.04 | 1175.89 | 97.68 | 0.00 | Protein coding |

Chrom, chromosome; Exp, expression; TPM, transcript per million; FDR, false discovery rate; NMD, nonsense mediated decay

Table S11: Top 100 differentially expressed transcript isoforms of RNA-binding factors, including RNA modifiers and splicing factors

| Name | ENSEMBL | Chrom | Exp in mGC (TPM) | Exp in cGC (TPM) | Fold change | FDR p-value | Biotype |
|---|---|---|---|---|---|---|---|
| *Ywhaz-204* | ENSMUST00000110362 | 15 | 362.99 | 0.01 | -32082.51 | 0.00 | Protein coding |
| *Psap-201* | ENSMUST00000105465 | 10 | 708.04 | 0.03 | -20793.65 | 0.00 | Protein coding |
| *Sidt2-201* | ENSMUST00000038488 | 9 | 429.27 | 0.02 | -19752.86 | 0.00 | Protein coding |
| *Fus-201* | ENSMUST00000077609 | 7 | 242.87 | 0.01 | -18584.48 | 0.00 | Protein coding |
| *Acly-201* | ENSMUST00000007131 | 11 | 243.28 | 0.02 | -14239.63 | 0.00 | Protein coding |
| *Uba52-208* | ENSMUST00000165126 | 8 | 781.91 | 0.06 | -12611.11 | 0.00 | Protein coding |
| *Rps10-208* | ENSMUST00000178774 | 17 | 492.25 | 0.05 | -10197.83 | 0.00 | Protein coding |
| *Rpl28-202* | ENSMUST00000078432 | 7 | 932.43 | 0.27 | -3488.23 | 0.00 | Protein coding |
| *Rpl7-205* | ENSMUST00000149566 | 1 | 870.57 | 0.61 | -1431.79 | 0.00 | Protein coding |
| *Rps10-201* | ENSMUST00000025052 | 17 | 555.07 | 0.86 | -643.90 | 0.00 | Protein coding |
| *Serpinh1-201* | ENSMUST00000094154 | 7 | 1066.74 | 1.70 | -626.96 | 0.00 | Protein coding |
| *Rpl18a-207* | ENSMUST00000212796 | 8 | 736.31 | 2.09 | -351.64 | 0.00 | Protein coding |
| *Rps7-203* | ENSMUST00000221871 | 12 | 1164.89 | 3.81 | -305.77 | 0.00 | Protein coding |
| *Rps14-205* | ENSMUST00000137400 | 18 | 1518.83 | 5.15 | -294.78 | 0.00 | Protein coding |
| *Rpl18a-208* | ENSMUST00000213053 | 8 | 405.94 | 1.88 | -216.21 | 0.00 | NMD |
| *Ctnnb1-201* | ENSMUST00000007130 | 9 | 385.25 | 2.43 | -158.35 | 0.00 | Protein coding |
| *Vcl-201* | ENSMUST00000022369 | 14 | 252.08 | 2.31 | -109.13 | 0.00 | Protein coding |
| *Ptbp1-216* | ENSMUST00000172282 | 10 | 286.46 | 3.02 | -94.97 | 0.00 | Protein coding |
| *Txndc5-201* | ENSMUST00000035988 | 13 | 416.72 | 4.46 | -93.40 | 0.00 | Protein coding |
| *Aldh2-202* | ENSMUST00000129753 | 5 | 254.20 | 2.93 | -86.65 | 0.00 | Protein coding |
| *Actg1-202* | ENSMUST00000071555 | 11 | 2549.80 | 29.50 | -86.42 | 0.00 | Protein coding |
| *Rps3-202* | ENSMUST00000107096 | 7 | 404.57 | 5.70 | -70.96 | 0.00 | Protein coding |
| *Lmnb1-201* | ENSMUST00000025486 | 18 | 503.51 | 8.15 | -61.79 | 0.00 | Protein coding |
| *Ctsd-202* | ENSMUST00000151120 | 7 | 364.57 | 5.98 | -60.98 | 0.00 | Protein coding |
| *Rpl19-202* | ENSMUST00000092425 | 11 | 292.07 | 5.12 | -56.99 | 0.00 | Protein coding |
| *Tpt1-201* | ENSMUST00000110894 | 14 | 3140.30 | 56.41 | -55.67 | 0.00 | Protein coding |
| *Acly-203* | ENSMUST00000107389 | 11 | 267.84 | 5.27 | -50.79 | 0.00 | Protein coding |

| | | | | | | | |
|---|---|---|---|---|---|---|---|
| *Calu-201* | ENSMUST00000031779 | 6 | 393.28 | 8.78 | -44.81 | 0.00 | Protein coding |
| *Atp1a1-201* | ENSMUST00000036493 | 3 | 527.52 | 11.87 | -44.43 | 0.00 | Protein coding |
| *Hnrnpa3-204* | ENSMUST00000111964 | 2 | 861.78 | 19.88 | -43.34 | 0.00 | Protein coding |
| *Rps26-202* | ENSMUST00000219591 | 10 | 1279.02 | 36.98 | -34.59 | 0.00 | retained_intron |
| *Junb-201* | ENSMUST00000064922 | 8 | 889.68 | 27.80 | -32.00 | 0.00 | Protein coding |
| *Rpl10a-212* | ENSMUST00000233427 | 17 | 449.57 | 16.13 | -27.87 | 0.00 | Protein coding |
| *Ncbp1-201* | ENSMUST00000030014 | 4 | 263.59 | 9.74 | -27.07 | 0.00 | Protein coding |
| *Rpsa-201* | ENSMUST00000035105 | 9 | 259.86 | 10.76 | -24.16 | 0.00 | Protein coding |
| *Hnrnpab-203* | ENSMUST00000109103 | 11 | 439.08 | 22.21 | -19.77 | 0.00 | Protein coding |
| *Rpl8-201* | ENSMUST00000004072 | 15 | 1493.97 | 76.61 | -19.50 | 0.00 | Protein coding |
| *Rpl37-202* | ENSMUST00000226758 | 15 | 502.14 | 25.98 | -19.33 | 0.00 | CDS not defined |
| *Eef1g-201* | ENSMUST00000052248 | 19 | 446.23 | 23.08 | -19.33 | 0.00 | Protein coding |
| *Mfge8-202* | ENSMUST00000107409 | 7 | 269.91 | 14.10 | -19.14 | 0.00 | Protein coding |
| *Rps15-202* | ENSMUST00000068408 | 10 | 1529.92 | 92.28 | -16.58 | 0.00 | Protein coding |
| *Hnrnpa2b1-208* | ENSMUST00000204158 | 6 | 330.38 | 20.15 | -16.40 | 0.00 | NMD |
| *Rcn3-201* | ENSMUST00000019683 | 7 | 311.86 | 19.07 | -16.35 | 0.00 | Protein coding |
| *Arhgdia-202* | ENSMUST00000106197 | 11 | 239.14 | 17.71 | -13.50 | 0.00 | Protein coding |
| *Sod1-201* | ENSMUST00000023707 | 16 | 890.64 | 66.12 | -13.47 | 0.00 | Protein coding |
| *Rpl35-203* | ENSMUST00000152441 | 2 | 275.80 | 23.31 | -11.83 | 0.00 | CDS not defined |
| *Vim-201* | ENSMUST00000028062 | 2 | 282.34 | 25.28 | -11.17 | 0.00 | Protein coding |
| *Rpl30-201* | ENSMUST00000009039 | 15 | 514.64 | 47.30 | -10.88 | 0.00 | Protein coding |
| *Slc25a3-201* | ENSMUST00000076694 | 10 | 1152.98 | 108.16 | -10.66 | 0.00 | Protein coding |
| *Rpl31-202* | ENSMUST00000178079 | 1 | 1178.03 | 111.66 | -10.55 | 0.00 | Protein coding |
| *Atp5pd-201* | ENSMUST00000043931 | 11 | 76.77 | 809.93 | 10.55 | 0.00 | Protein coding |
| *Psma7-201* | ENSMUST00000029082 | 2 | 77.42 | 919.00 | 11.87 | 0.00 | Protein coding |
| *Rps14-201* | ENSMUST00000025511 | 18 | 246.90 | 2955.36 | 11.97 | 0.00 | Protein coding |
| *Eif3e-201* | ENSMUST00000022960 | 15 | 78.86 | 962.08 | 12.20 | 0.00 | Protein coding |
| *Prdx4-201* | ENSMUST00000026328 | X | 83.77 | 1024.49 | 12.23 | 0.00 | Protein coding |
| *Naca-201* | ENSMUST00000073868 | 10 | 251.19 | 3323.30 | 13.23 | 0.00 | Protein coding |
| *Calm1-202* | ENSMUST00000110082 | 12 | 77.28 | 1027.87 | 13.30 | 0.00 | Protein coding |
| *Sub1-202* | ENSMUST00000110504 | 15 | 65.15 | 939.43 | 14.42 | 0.00 | Protein coding |

| | | | | | | | |
|---|---|---|---|---|---|---|---|
| *Eif3g-201* | ENSMUST00000004206 | 9 | 52.13 | 767.83 | 14.73 | 0.00 | Protein coding |
| *Skp1-201* | ENSMUST00000037324 | 11 | 95.87 | 1412.21 | 14.73 | 0.00 | Protein coding |
| *Ranbp1-202* | ENSMUST00000115645 | 16 | 69.45 | 1097.25 | 15.80 | 0.00 | Protein coding |
| *Ywhaq-201* | ENSMUST00000103002 | 12 | 86.40 | 1677.83 | 19.42 | 0.00 | Protein coding |
| *Fabp5-201* | ENSMUST00000029046 | 3 | 40.22 | 787.59 | 19.58 | 0.00 | Protein coding |
| *Vdac3-201* | ENSMUST00000009036 | 8 | 42.73 | 895.09 | 20.95 | 0.00 | Protein coding |
| *Cnbp-203* | ENSMUST00000113619 | 6 | 83.34 | 1757.71 | 21.09 | 0.00 | Protein coding |
| *Hsd17b10-201* | ENSMUST00000026289 | X | 47.64 | 1016.55 | 21.34 | 0.00 | Protein coding |
| *Mdh1-206* | ENSMUST00000239073 | 11 | 73.61 | 1578.96 | 21.45 | 0.00 | Protein coding |
| *Atp5f1c-203* | ENSMUST00000114897 | 2 | 74.92 | 1698.51 | 22.67 | 0.00 | Protein coding |
| *Hsp90aa1-201* | ENSMUST00000021698 | 12 | 51.65 | 1203.55 | 23.30 | 0.00 | Protein coding |
| *Hmgb2-201* | ENSMUST00000067925 | 8 | 47.94 | 1134.21 | 23.66 | 0.00 | Protein coding |
| *Bud31-207* | ENSMUST00000162594 | 5 | 33.40 | 879.74 | 26.34 | 0.00 | Protein coding |
| *Rps27l-201* | ENSMUST00000040917 | 9 | 74.03 | 2035.22 | 27.49 | 0.00 | Protein coding |
| *Eloc-201* | ENSMUST00000115352 | 1 | 27.30 | 874.13 | 32.02 | 0.00 | Protein coding |
| *Polr2g-201* | ENSMUST00000096261 | 19 | 21.06 | 739.49 | 35.11 | 0.00 | Protein coding |
| *Manf-201* | ENSMUST00000069036 | 9 | 71.09 | 2608.33 | 36.69 | 0.00 | Protein coding |
| *Rps27a-202* | ENSMUST00000102845 | 11 | 21.80 | 930.06 | 42.67 | 0.00 | Protein coding |
| *Sumo1-201* | ENSMUST00000091374 | 1 | 42.60 | 2211.55 | 51.91 | 0.00 | Protein coding |
| *Bnip3-201* | ENSMUST00000106112 | 7 | 19.52 | 1113.90 | 57.07 | 0.00 | Protein coding |
| *Rbbp7-202* | ENSMUST00000112326 | X | 16.76 | 1159.36 | 69.16 | 0.00 | Protein coding |
| *Rpf2-206* | ENSMUST00000183309 | 10 | 9.86 | 754.24 | 76.48 | 0.00 | Protein coding |
| *Rpl10-201* | ENSMUST00000008826 | X | 12.37 | 1061.29 | 85.83 | 0.00 | Protein coding |
| *Hspa8-202* | ENSMUST00000117557 | 9 | 12.04 | 1175.89 | 97.68 | 0.00 | Protein coding |
| *Srsf7-208* | ENSMUST00000235069 | 17 | 16.64 | 1828.31 | 109.86 | 0.00 | Protein coding |
| *Prdx1-202* | ENSMUST00000106470 | 4 | 57.15 | 7560.73 | 132.30 | 0.00 | Protein coding |
| *Atp5f1b-205* | ENSMUST00000126040 | 10 | 4.60 | 1206.92 | 262.42 | 0.00 | Retained intron |
| *Myef2-207* | ENSMUST00000147105 | 2 | 3.31 | 960.85 | 290.41 | 0.00 | Protein coding |
| *Ubb-202* | ENSMUST00000136938 | 11 | 4.29 | 1298.06 | 302.44 | 0.00 | Protein coding |
| *Gstp1-208* | ENSMUST00000237893 | 19 | 1.03 | 1048.34 | 1016.24 | 0.00 | Protein coding |
| *Rplp0-204* | ENSMUST00000156359 | 5 | 1.46 | 4475.28 | 3070.70 | 0.00 | Protein coding |

| | | | | | | | |
|---|---|---|---|---|---|---|---|
| *Tpt1-204* | ENSMUST00000134040 | 14 | 0.24 | 775.38 | 3184.26 | 0.00 | CDS not defined |
| *Rpl14-205* | ENSMUST00000216990 | 9 | 0.16 | 1096.43 | 6917.52 | 0.00 | NMD |
| *Btf3-204* | ENSMUST00000134542 | 13 | 0.21 | 1509.03 | 7292.53 | 0.00 | Protein coding |
| *Ppia-201* | ENSMUST00000090749 | 11 | 1.62 | 23437.87 | 14440.09 | 0.00 | NMD |
| *Rack1-202* | ENSMUST00000125166 | 11 | 0.03 | 803.69 | 25082.95 | 0.00 | CDS not defined |
| *Hsp90aa1-208* | ENSMUST00000155242 | 12 | 0.04 | 1633.93 | 39758.74 | 0.00 | Protein coding |
| *Ranbp1-201* | ENSMUST00000052325 | 16 | 0.04 | 2799.31 | 65768.81 | 0.00 | Protein coding |
| *Fkbp3-205* | ENSMUST00000221166 | 12 | 0.01 | 1053.54 | 134735.85 | 0.00 | NMD |
| *Eif3m-202* | ENSMUST00000111110 | 2 | 0.01 | 1506.49 | 212546.89 | 0.00 | Protein coding |
| *Actg1-204* | ENSMUST00000106215 | 11 | 0.00 | 948.29 | 317548.00 | 0.00 | Protein coding |
| *Prdx2-208* | ENSMUST00000164807 | 8 | 0.00 | 1684.12 | 370134.20 | 0.00 | Protein coding |

Chrom, chromosome; Exp, expression; TPM, transcript per million; FDR, false discovery rate; NMD, nonsense mediated decay

Table S12: Top 50 differentially expressed transcript isoforms of transcription terminators and polyadenylation factors

| *Name* | ENSEMBL | Chrom | Exp in mGC (TPM) | Exp in cGC (TPM) | Fold change | FDR p-value | Biotype |
|---|---|---|---|---|---|---|---|
| *Rai1-201* | ENSMUST00000064190 | 11 | 22.10 | 0.00 | -6745.83 | 0.00 | Protein coding |
| *Scaf4-201* | ENSMUST00000039280 | 16 | 17.47 | 0.01 | -3095.38 | 0.00 | Protein coding |
| *Sympk-201* | ENSMUST00000023882 | 7 | 48.98 | 0.02 | -2384.02 | 0.00 | Protein coding |
| *Pabpc4-203* | ENSMUST00000106241 | 4 | 14.96 | 0.01 | -1830.64 | 0.00 | Protein coding |
| *Rai1-203* | ENSMUST00000102688 | 11 | 8.81 | 0.01 | -1583.05 | 0.00 | Protein coding |
| *Pabpn1-201* | ENSMUST00000022808 | 14 | 128.79 | 0.13 | -1027.12 | 0.00 | Protein coding |
| *Pabpn1-207* | ENSMUST00000150975 | 14 | 54.40 | 0.07 | -821.37 | 0.00 | Protein coding |
| *Srsf1-206* | ENSMUST00000171976 | 11 | 23.91 | 0.04 | -642.78 | 0.00 | Retained intron |
| *Cdk9-208* | ENSMUST00000154131 | 2 | 15.80 | 0.03 | -556.80 | 0.00 | NMD |
| *Rai1-206* | ENSMUST00000156902 | 11 | 11.10 | 0.03 | -326.86 | 0.00 | CDS not defined |
| *Elavl1-203* | ENSMUST00000208987 | 8 | 11.74 | 0.04 | -293.56 | 0.00 | Retained intron |
| *Rbm3-210* | ENSMUST00000145450 | X | 13.52 | 0.12 | -115.24 | 0.00 | Retained intron |
| *Pabpn1-203* | ENSMUST00000139985 | 14 | 13.90 | 0.15 | -90.41 | 0.00 | Protein coding |
| *Srsf1-202* | ENSMUST00000107920 | 11 | 12.16 | 0.17 | -72.64 | 0.00 | Protein coding |
| *Sympk-206* | ENSMUST00000146903 | 7 | 14.86 | 0.27 | -54.59 | 0.00 | NMD |
| *Pcf11-205* | ENSMUST00000208255 | 7 | 12.02 | 0.26 | -46.06 | 0.00 | Protein coding |
| *Pabpc4-208* | ENSMUST00000183940 | 4 | 16.05 | 0.38 | -41.90 | 0.00 | NMD |
| *Pabpn1-206* | ENSMUST00000146271 | 14 | 9.23 | 0.29 | -32.37 | 0.00 | Retained intron |
| *Papola-215* | ENSMUST00000170002 | 12 | 13.27 | 0.54 | -24.49 | 0.00 | Protein coding |
| *Cdk9-201* | ENSMUST00000009699 | 2 | 55.15 | 2.84 | -19.43 | 0.00 | Protein coding |
| *Pabpc4-204* | ENSMUST00000106243 | 4 | 33.71 | 2.38 | -14.15 | 0.00 | Protein coding |
| *Pabpc1-205* | ENSMUST00000147116 | 15 | 9.26 | 0.66 | -14.10 | 0.00 | Retained intron |
| *Srsf7-207* | ENSMUST00000235036 | 17 | 83.49 | 7.27 | -11.49 | 0.00 | Retained intron |
| *Sf3b1-202* | ENSMUST00000185429 | 1 | 9.33 | 0.82 | -11.39 | 0.00 | Retained intron |
| *Fip1l1-203* | ENSMUST00000113535 | 5 | 30.43 | 3.33 | -9.15 | 0.00 | Protein coding |
| *Pabpc1-202* | ENSMUST00000142357 | 15 | 9.89 | 1.39 | -7.13 | 0.00 | Retained intron |
| *Pabpc4-202* | ENSMUST00000080178 | 4 | 72.02 | 10.67 | -6.75 | 0.00 | Protein coding |
| *Srsf7-204* | ENSMUST00000234577 | 17 | 9.63 | 1.45 | -6.63 | 0.00 | Retained intron |

| | | | | | | | |
|---|---|---|---|---|---|---|---|
| *Pabpc1-201* | ENSMUST00000001809 | 15 | 715.81 | 124.71 | -5.74 | 0.00 | Protein coding |
| *Srsf1-203* | ENSMUST00000132983 | 11 | 27.33 | 5.31 | -5.15 | 0.00 | Retained intron |
| *Rbm3-202* | ENSMUST00000115615 | X | 31.03 | 199.55 | 6.43 | 0.00 | Protein coding |
| *Rbm3-204* | ENSMUST00000115617 | X | 19.52 | 130.18 | 6.67 | 0.00 | Protein coding |
| *Rbm3-203* | ENSMUST00000115616 | X | 45.84 | 348.38 | 7.60 | 0.00 | Protein coding |
| *Pabpn1-208* | ENSMUST00000172557 | 14 | 2.50 | 22.21 | 8.90 | 0.00 | Protein coding |
| *Srsf7-206* | ENSMUST00000234889 | 17 | 10.10 | 92.75 | 9.18 | 0.00 | Retained intron |
| *Rbm3-206* | ENSMUST00000115621 | X | 30.20 | 280.28 | 9.28 | 0.00 | Protein coding |
| *Srsf7-203* | ENSMUST00000234503 | 17 | 8.58 | 97.13 | 11.32 | 0.00 | NMD |
| *Nudt21-204* | ENSMUST00000212981 | 8 | 21.24 | 257.82 | 12.14 | 0.00 | Protein coding |
| *Pin1-201* | ENSMUST00000034689 | 9 | 6.74 | 84.17 | 12.49 | 0.00 | Protein coding |
| *Srsf1-204* | ENSMUST00000134824 | 11 | 8.12 | 113.25 | 13.94 | 0.00 | NMD |
| *Ssu72-201* | ENSMUST00000030905 | 4 | 14.37 | 219.29 | 15.26 | 0.00 | Protein coding |
| *Pabpc4-206* | ENSMUST00000146156 | 4 | 2.59 | 42.66 | 16.50 | 0.00 | Retained intron |
| *Srsf7-201* | ENSMUST00000063417 | 17 | 27.02 | 621.28 | 22.99 | 0.00 | Protein coding |
| *Wdr33-205* | ENSMUST00000234344 | 18 | 2.76 | 63.61 | 23.04 | 0.00 | Protein coding |
| *Wdr33-202* | ENSMUST00000082319 | 18 | 1.81 | 51.42 | 28.41 | 0.00 | Protein coding |
| *Srsf7-205* | ENSMUST00000234696 | 17 | 9.08 | 338.39 | 37.25 | 0.00 | Protein coding |
| *Clp1-202* | ENSMUST00000138231 | 2 | 0.41 | 22.15 | 54.16 | 0.00 | CDS not defined |
| *Papola-219* | ENSMUST00000172040 | 12 | 0.81 | 47.64 | 58.70 | 0.00 | CDS not defined |
| *Pabpc1-204* | ENSMUST00000146577 | 15 | 2.39 | 254.57 | 106.37 | 0.00 | Retained intron |
| *Srsf7-208* | ENSMUST00000235069 | 17 | 16.64 | 1828.31 | 109.86 | 0.00 | Protein coding |

Chrom, chromosome; Exp, expression; TPM, transcript per million; FDR, false discovery rate; NMD, nonsense mediated decay
